# The IRE1α/XBP1 signaling axis drives myoblast fusion in adult skeletal muscle

**DOI:** 10.1101/2023.09.26.559625

**Authors:** Aniket S. Joshi, Meiricris Tomaz da Silva, Anirban Roy, Tatiana E. Koike, Mingfu Wu, Micah B. Castillo, Preethi H. Gunaratne, Yu Liu, Takao Iwawaki, Ashok Kumar

## Abstract

Skeletal muscle regeneration involves a signaling network that regulates the proliferation, differentiation, and fusion of muscle precursor cells to injured myofibers. Inositol requiring enzyme 1 alpha (IRE1α) is one of the arms of the unfolded protein response (UPR) that regulates cellular proteostasis in response to ER stress. Here, we demonstrate that inducible deletion of IRE1α in adult muscle stem cells (i.e. satellite cells) of mice impairs skeletal muscle regeneration primarily through inhibiting myoblast fusion step. Knockdown of IRE1α or its downstream target, X-box protein 1 (XBP1), also inhibits fusion of cultured myoblasts during myogenesis. Genome-wide transcriptome analysis revealed that knockdown of IRE1α or XBP1 deregulates the gene expression of molecules involved in the regulation of myoblast fusion. The IRE1α-XBP1 axis mediates the gene expression of multiple profusion molecules, including Myomaker (*Mymk*) during myogenic differentiation. Our study demonstrates that spliced XBP1 (sXBP1) transcription factor binds to the promoter region of *Mymk* gene during myogenesis. Overexpression of myomaker in IRE1α-knockdown cultures rescues fusion defects. Finally, our results show that inducible deletion of *IRE1*α in satellite cells inhibits myoblast fusion and myofiber hypertrophy in response to functional overload. Collectively, our study demonstrates that IRE1α promotes myoblast fusion through sXBP1-mediated up-regulation in the gene expression of profusion molecules.

**Significance Statement:** Myoblast fusion is an essential step for regeneration and post-natal growth of skeletal muscle. We demonstrate that the activation of the IRE1α/XBP1 arm of the unfolded protein response induces myoblast fusion through augmenting the gene expression of multiple profusion molecules, including myomaker. This study has identified a novel signaling axis that link ER stress-induced non-myogenic signaling pathway to myoblast fusion. Augmenting the activity of IRE1α/XBP1 pathway could be a potential therapeutic strategy for various muscle degenerative diseases.

## INTRODUCTION

Skeletal muscle cells, more commonly known as myofibers, are multinucleated syncytia that arise by the fusion of thousands of myoblasts during development (1). Adult skeletal muscle tissue retains regenerative capability mainly due to the presence of a pool of muscle progenitor cells, known as satellite cells, which reside in basal lamina around myofibers in a quiescent state (2). Following muscle injury, satellite cells are activated, which then proliferate, differentiate into myoblasts, and form multinucleated myotubes through the fusion of myoblast with another myoblast or nascent myotubes. Moreover, a subset of myoblasts also fuses with damaged pre-existing myofibers to accomplish muscle repair (2, 3).

Myoblast fusion is a systematic process that involves migration and alignment of membranes of fusion-competent myoblasts, remodelling of cytoskeleton at contact sites followed by opening of fusion pores to allow movement of cytoplasmic content, and eventually amalgamation of two myogenic cells into one (4–6). Recently, two transmembrane proteins, named Myomaker and Myomerger (also known as Myomixer and Minion) have been identified as major drivers of myoblast fusion in diverse conditions (7–10). It is also now increasingly clear that myoblast fusion is regulated by multiple signaling pathways that are activated due to interaction of specific membrane proteins between fusion partners or as a part of myogenic differentiation program (11). However, the molecular and signaling mechanisms regulating myoblast fusion remain less understood.

Skeletal muscle regeneration requires synthesis and processing of a large number of proteins, including growth factors, signaling proteins, and a new set of contractile, cytoskeletal, and membrane proteins (2). The endoplasmic reticulum (ER) is the site for folding of membrane and secretory proteins, synthesis of lipids and sterols, and storage of free calcium. Physiological demands or pathological stresses, such as the presence of mutated proteins that cannot properly fold in the ER, can lead to accumulation of unfolded protein, thereby causing ER stress (12). Mammalian cells respond to the presence of unfolded or misfolded proteins within the ER through the activation of an intricate set of signal pathways, collectively termed as the unfolded protein response (UPR) that realigns ER folding capacity and restores cellular homeostasis. The UPR consist of three ER-resident proteins: inositol requiring enzyme 1α/β (IRE1), PKR-like ER kinase (PERK), and activating transcription factor 6α/β (ATF6) (13, 14). Among these, the IRE1 is the most ancient and conserved branch of the UPR that plays a major role in resolving ER stress. The cytosolic portion of IRE1 possesses a kinase domain that autophosphorylates, leading to the stimulation of its endoribonuclease activity, which catalyzes unconventional processing of the mRNA encoding X-box-binding protein 1 (XBP1) and creates a transcriptionally active XBP1, known as spliced XBP1 (sXBP1). This results in the stimulation of gene expression of many molecules that increase the protein folding capacity as well as augment protein degradation and transport pathways, which mitigates the burden of misfolded protein within the ER. IRE1 activation can also lead to promiscuous endoribonuclease activity that causes mRNA decay at the ER membrane, a process called Regulated IRE1-Dependent Decay (RIDD), thus reducing the protein load. Finally, during ER stress, the kinase domain of IRE1 binds to the TRAF2 adaptor protein that results in the phosphorylation and activation of JNK to control cell fate (13–15). Remarkably, signaling pathways activated by stimulation of various cell surface receptors have been found to crosstalk with UPR signaling in an ER stress-independent manner suggesting that ER sensors also mediate non-canonical UPR responses to regulate cell physiology (16, 17).

Accumulating evidence suggests that UPR pathways play important roles in skeletal muscle development and regenerative myogenesis (18, 19). Pharmacological activation of ER stress has been found to augment myotube formation in cultured myoblasts following induction of differentiation (20). Recent studies have also demonstrated that the PERK arm of the UPR is essential for self-renewal of satellite cells (21, 22). We have recently demonstrated that IRE1α signaling in myofibers promotes skeletal muscle regeneration following acute injury as well as in the mdx model of Duchenne muscular dystrophy (23). However, the cell autonomous role of IRE1α in the regulation of satellite cell function in adult skeletal muscle remained completely unknown.

In the present study, using genetic and molecular approaches, we demonstrate that satellite cell-specific ablation of IRE1α impairs skeletal muscle regeneration in adult mice. Importantly, IRE1α does not influence satellite cell abundance or differentiation in regenerating skeletal muscle. Rather, IRE1α promotes muscle repair and growth through augmenting myoblast fusion. Overexpression of IRE1α or its downstream target sXBP1 in cultured myoblasts results in the formation of myotubes having an increased diameter following induction of differentiation. The IRE1α/XBP1 signaling augments the gene expression of multiple profusion molecules, including myomaker in differentiating myoblasts. Finally, our results demonstrate that IRE1α signaling in muscle progenitor cells is also essential for overload-induced myofiber hypertrophy in adult mice.

## RESULTS

### Activation of IRE1α in muscle progenitor cells

Pax7 transcription factor is a widely used marker expressed in both quiescent and activated satellite cells (24). By performing immunofluorescence assay, we first examined whether IRE1α protein is activated in satellite cells. TA muscle of wild-type mice was injured by intramuscular injection of 1.2% BaCl_2_ solution. Results showed that p-IRE1α co-localized with Pax7 protein in both uninjured and injured TA muscle, suggesting that IRE1α is activated in both quiescent and activated satellite cells *in vivo* (**Fig. 1A**). We next studied the activation of IRE1α in cultured myogenic cells. Mouse primary myoblasts incubated in growth medium (GM) were immunostained for Pax7, and p-IRE1α or total IRE1α protein. Consistent with *in vivo* results, we found that both p-IRE1α and IRE1α protein were present in Pax7^+^ cells **(Fig. 1B)**. Following a FACS-based intracellular protein detection assay, we also investigated whether p-IRE1α protein is present in cultured muscle progenitor cells. Mouse primary myoblasts incubated in GM were collected and analysed by FACS for α7-integrin and p-IRE1α or IRE1α protein. Results showed that p-IRE1α and total IRE1α protein were present in the α7-integrin^+^ satellite cells **(Fig. 1C)**. We next sought to determine how the levels of p-IRE1α and total IRE1α protein are regulated during myogenic differentiation. Primary myoblasts were incubated in differentiation medium (DM) for 0, 6, 12, 24, 48 and 72 h and the cell extracts made were analysed by western blot. Results showed that p-IRE1α and total IRE1α proteins along with Pax7 protein were highly abundant in myoblast cultures incubated in GM. However, the levels of p-IRE1α, IRE1α, and Pax7 protein were gradually reduced whereas the muscle differentiation markers, myogenin and myosin heavy chain (MyHC) were increased after incubation in DM **(Fig. 1D)**. In response to ER stress, IRE1α gets activated which causes an unconventional splicing of XBP1 mRNA resulting in the formation of spliced XBP1 (sXBP1) mRNA (25). Western blot analysis showed a peak increase in the protein levels of sXBP1 at 24 h of addition of DM **(Fig. 1D)**. These results suggest that the IRE1α/XBP1 arm of the UPR is activated in muscle progenitor cells both *in vivo* and *in vitro*.

**FIGURE 1.**
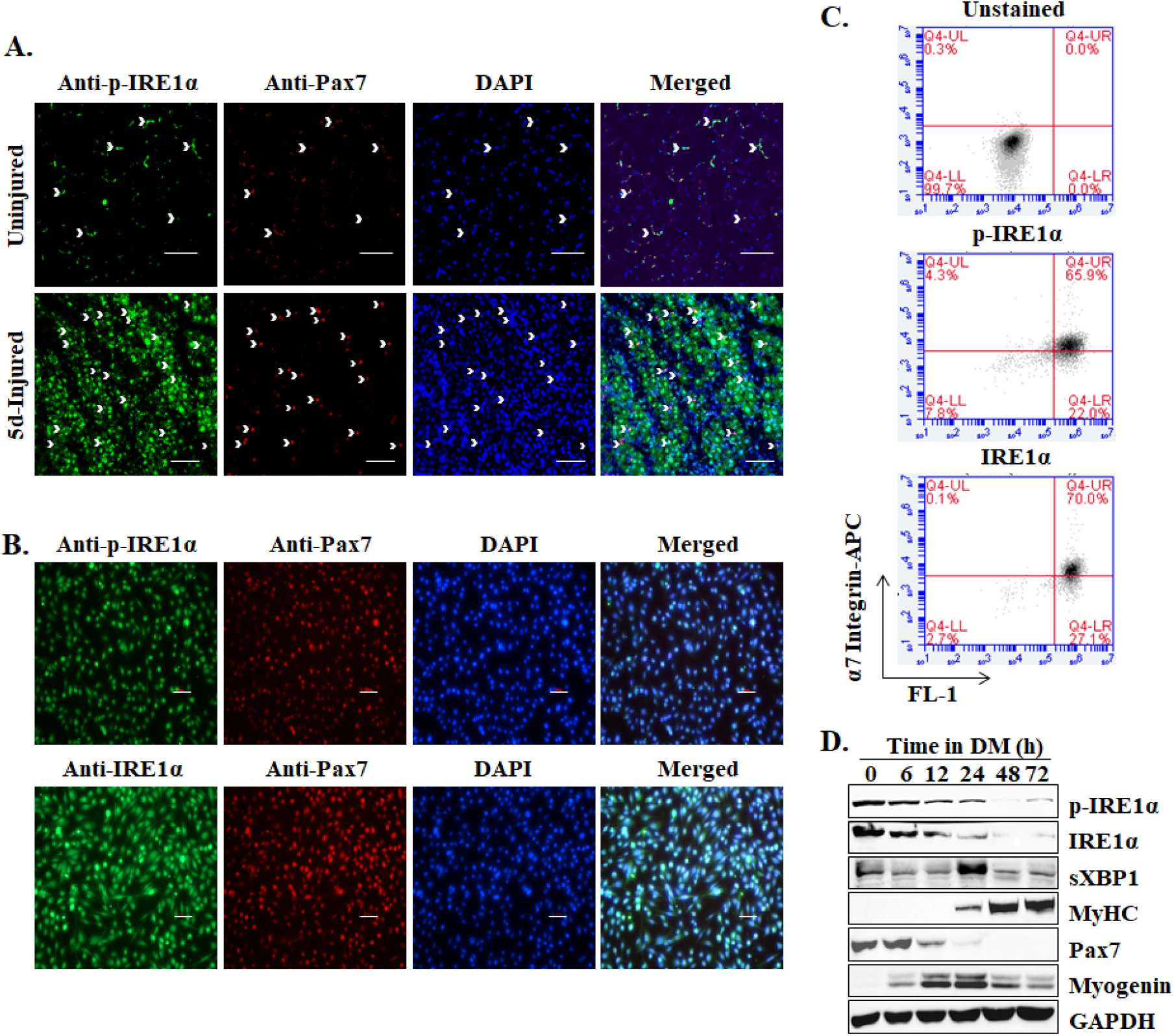
Activation of IRE1α is muscle progenitor cells. **(A)** Uninjured and 5d-injured TA muscle sections of wild-type mice were immunostained for phosho-IRE1 (p-IRE1) and Pax7 protein. Nuclei were counterstained with DAPI. Representative photomicrographs presented here demonstrate that p-IRE1 is present in Pax7^+^ cells in both uninjured and injured muscle. Scale bar: 100 µm. **(B)** Primary myoblasts prepared from wild-type mice were fixed and immunostained for p-IRE1α or IRE1α and Pax7 protein. DAPI was used to counterstain nuclei. Representative photomicrographs are presented here. Scale bar: 50 µm. **(C)** Cultured myoblasts were subjected for FACS-based analysis for the expression of α7β1-integrin and p-IRE1 or total IRE1α. Representative scatter plots demonstrate presence of phospho-IRE1^+^ and IRE1^+^ cells amongst α7β1-intigrin^+^ population. n=3 in each group. (**D)** Cultured myoblasts were incubated in DM for indicated time points and analyzed by performing western blot. Immunoblots present here demonstrate the relative levels of p-IRE1α, IRE1α, sXBP1, MyHC, Pax7, myogenin, and an unrelated protein GAPDH.

### Ablation of IRE1α in satellite cells inhibits skeletal muscle regeneration

We next investigated whether IRE1α signaling in satellite cells affects skeletal muscle regeneration in adult mice. Floxed *Ern1* mice (henceforth *Ern1^fl/fl^*) were crossed with tamoxifen inducible satellite cell-specific Cre mice (*Pax7-CreERT2*) to generate *Pax7-CreERT2;Ern1^fl/fl^* (henceforth, *Ern1^scKO^*) and littermate *Ern1^fl/fl^* mice **(Fig. 2A)**. 8-week-old *Ern1^scKO^*mice were given intraperitoneal injection of tamoxifen for 4 consecutive days. The mice were maintained on tamoxifen-containing chow for the entire duration of the study to ensure IRE1α deletion. *Ern1^fl/fl^* mice were also subjected to the same tamoxifen regimen and served as controls. One side TA muscle was injured by intramuscular injection of 1.2 % BaCl_2_ solution whereas the contralateral muscle served as uninjured control. The TA muscle was isolated at day 5 or 14 post-injury **(Fig. 2B)**. Our analysis showed that wet weight of 5d-injured TA muscle normalized by body weight of *Ern1^scKO^* mice was significantly reduced compared to *Ern1^fl/fl^* mice **(Fig. 2C-E)**. Next, transverse sections of uninjured and injured TA muscle were generated analyzed by performing H&E staining **(Fig. 2F)**. Interestingly, average myofiber cross-sectional area (CSA) was significantly reduced in injured TA muscle of *Ern1^scKO^* mice compared to corresponding injured TA muscle of *Ern1^fl/fl^* mice both at 5d and 14d post-injury **(Fig. 2F-H)**. These results suggest that satellite cell-specific deletion of IRE1α inhibits muscle regeneration in adult mice.

**FIGURE 2.**
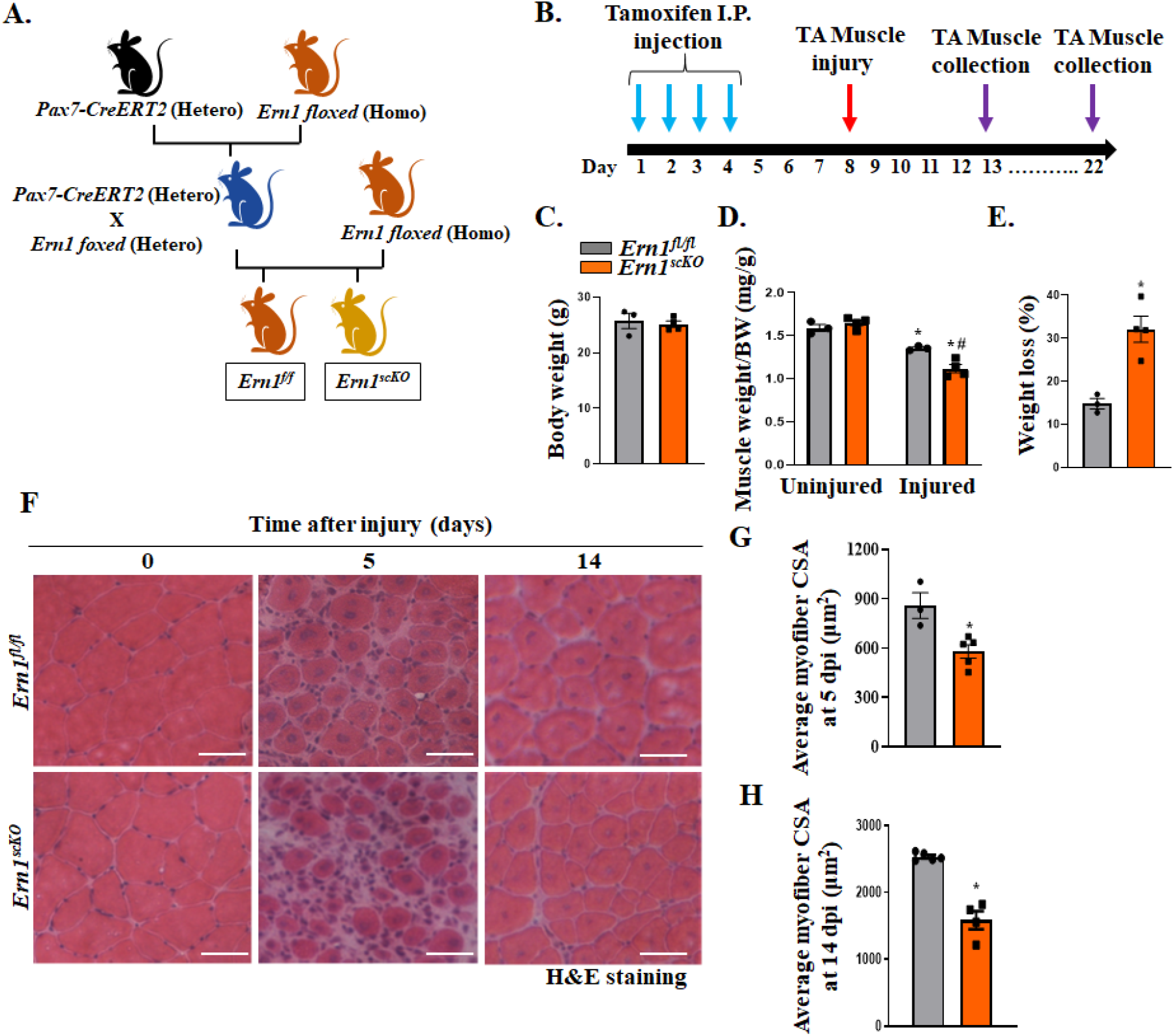
Satellite cell-specific ablation of IRE1α impairs skeletal muscle regeneration. **(A)** Schematic representation of breeding strategy used for the generation of *Ern1^fl/fl^* and *Ern1^scKO^* mice. **(B)** Schematic representation of the experimental design indicating time of Ern1 deletion, muscle injury, and collection. One side TA muscle of *Ern1^fl/fl^* and *Ern1^scKO^* mice was injured by intramuscular injection of BaCl_2_ whereas contralateral muscle served as control. At day 5 post-injury, the mice were analyzed for **(C)** overall body weight; **(D)** TA muscle wet weight normalized by body weight; and **(E)** loss in wet weight of injured TA muscle compared to contralateral uninjured muscle in percentage. Transverse sections of uninjured (day 0) and injured (days 5 or 14 after injury) TA muscle were generated and used for morphometric analysis. **(F)** Representative photomicrographs of Hematoxylin and Eosin (H&E) stained transverse TA muscle sections of *Ern1^fl/fl^* and *Ern1^scKO^* mice. Scale bar: 50µm. Quantitative analysis of average myofiber cross-sectional area (CSA) in TA muscle of *Ern1^fl/fl^* and *Ern1^scKO^* mice at **(G)** day 5 and **(H)** day 14 days post-injury. n=3-5 in each group. Data are presented as mean ± SEM and analyzed by unpaired Student *t* test or two-way ANOVA followed by Tukey’s multiple comparison test. *p ≤ 0.05; values significantly different from corresponding uninjured muscle; #p ≤ 0.05; values significantly different from injured TA muscle of *Ern1^fl/fl^* mice.

### Targeted ablation of IRE1α inhibits growth of newly formed myofibers

Skeletal muscle regeneration is dependent on the hierarchical expression of myogenic regulatory factors (MRFs, i.e. Myf5, MyoD, and myogenin) and the early regeneration marker, embryonic isoform of myosin-heavy chain (eMyHC), which ultimately determines the efficiency of muscle repair (26, 27). We next investigated the effect of IRE1α deletion in satellite cells on the level Transverse sections of uninjured and 5d-injured TA muscle of *Ern1^fl/fl^* and *Ern1^scKO^* mice were immunostained for eMyHC and laminin (to label boundaries of myofibers) protein. While there was no significant difference in the percentage of eMyHC^+^ myofibers per laminin-stained cells, a significant increase in the number of eMyHC^+^ cells per field was observed in 5d-injured TA muscle of *Ern1^scKO^* mice compared to corresponding muscle of *Ern1^fl/fl^* mice **(Fig. 3A-C)**. Moreover, average CSA and percentage of eMyHC^+^ myofibers containing 2 or more centrally located nuclei were significantly reduced in 5d-injured TA muscle of *Ern1^scKO^* mice compared to *Ern1^fl/fl^* mice **(Fig. 3D, E)**. Western blot analysis showed that protein levels of MyoD and myogenin were comparable whereas the levels of IRE1α protein were significantly reduced in 5d-injured TA muscle of *Ern1^scKO^* mice compared with corresponding 5d-injured TA muscle of *Ern1^fl/fl^* mice **(Fig. 3F, G)**. These results are consistent with our previously published report demonstrating that satellite cell-specific ablation of XBP1 (a downstream target of IRE1α) in adult mice does not affect the expression of MyoD, myogenin, or eMyHC in TA muscle at day 5 post-injury (21). We also investigated whether knockdown of IRE1α affects the abundance of myogenic regulatory factors (MRFs) in cultured primary myoblasts. Primary myoblasts were transfected with control or IRE1α siRNA for 24 h. The cells were then incubated in DM and collected at different time points after incubation in DM. There was no difference in the protein levels of myogenin between control and IRE1α knockdown cultures. However, a small reduction in levels of MyHC protein was observed in IRE1α knockdown cultures compared to corresponding controls at different time points after incubation in DM **(Fig. 3H)**.

**FIGURE 3.**
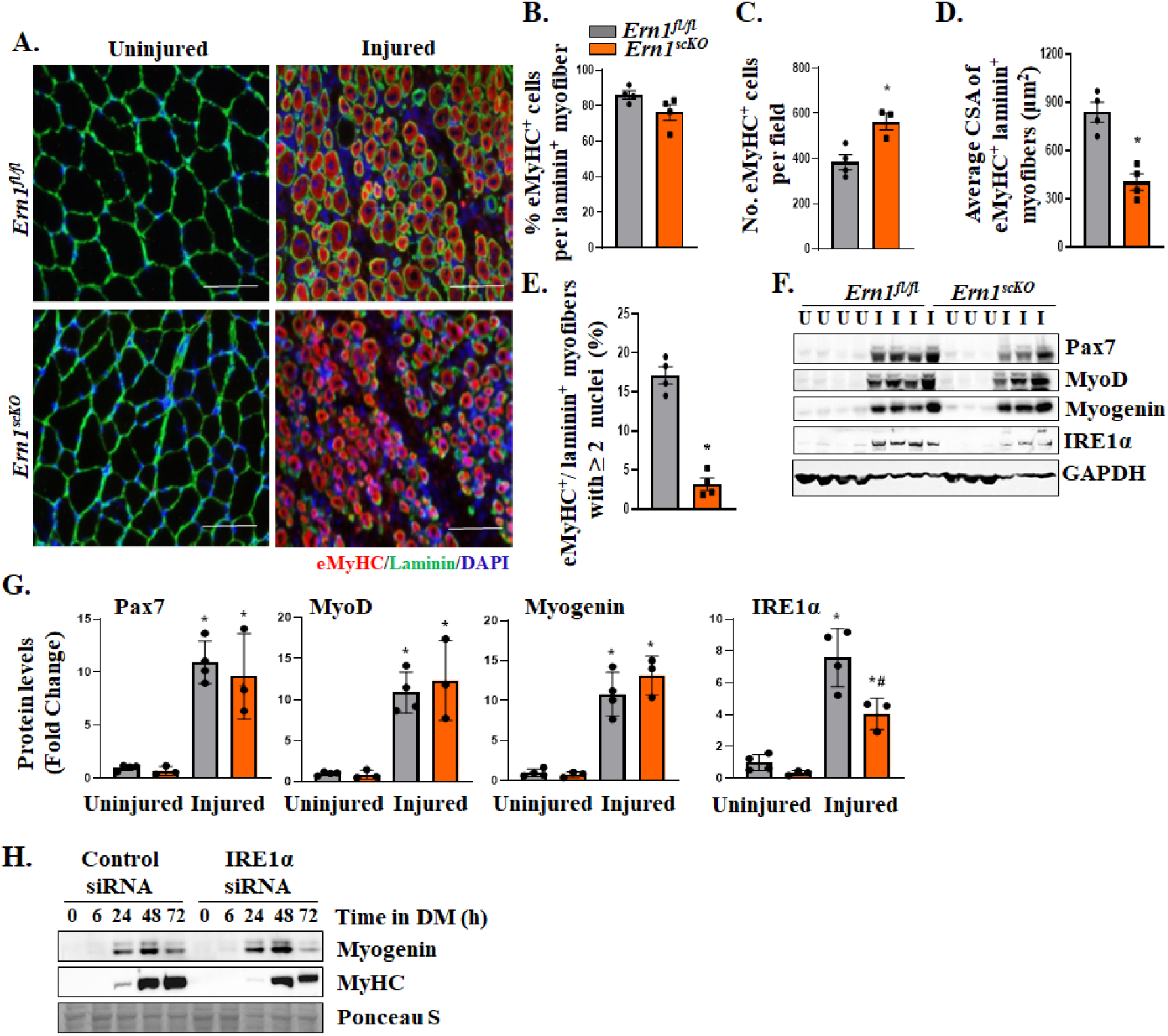
Loss of IRE1α in satellite cells inhibits formation of regenerating myofibers. **(A)** Representative photomicrographs of transverse sections of uninjured and 5d-injured TA muscle of *Ern1^fl/fl^* and *Ern1^scKO^* mice immunostained for embryonic isoform of myosin heavy chain (eMyHC) and laminin protein. DAPI was used to counterstain nuclei. Scale bar: 50µm. Quantification of **(B)** percentage of eMyHC^+^ cells per laminin^+^ myofibers, **(C)** number of eMyHC^+^ cells per field, **(D)** average CSA of eMyHC^+^ laminin^+^ myofibers, and **(E)** percentage of eMyHC^+^ laminin^+^ myofibers containing 2 or more nuclei. n=3-4 per group. Data are presented as mean ± SEM. *p ≤ 0.05, values significantly different from corresponding muscle of *Ern1^fl/fl^* mice analyzed by unpaired Student *t* test. **(F)** Western blot and (**G**) densitometry analysis showing protein levels of Pax7, MyoD, Myogenin, IRE1α, and an unrelated protein GAPDH in uninjured and 5d-injured TA muscle of *Ern1^fl/fl^* and *Ern1^scKO^* mice. n=3-4 in each group. *p ≤ 0.05, values significantly different from corresponding uninjured muscle analyzed by unpaired Student *t* test. #p ≤ 0.05, values significantly different from injured TA muscle of *Ern1^fl/fl^* mice analyzed by unpaired Student *t* test. **(H)** Primary myoblasts were treated with control or IRE1α siRNA for 24 h, followed by incubation in DM for indicated time points. Representative immunoblots presented here show the levels of Myogenin and MyHC protein at indicated time points after incubation in DM.

We next sought to determine whether targeted ablation of IRE1α affects satellite cell number in injured skeletal muscle of mice. There was no significant difference in the number of Pax7^+^ cells in 5d-injured TA muscle of *Ern1^fl/fl^* and *Ern1^scKO^* mice **(Fig. S1A, B)**. Additionally, there was no significant difference in protein levels of Pax7 in uninjured or 5d-injured TA muscle of *Ern1^fl/fl^* and *Ern1^scKO^* mice (**Fig. 3F, G**). We also investigated whether siRNA-mediated knockdown of IRE1α affects the number of satellite cells in cultured myogenic cells. There was no significant difference in the proportion of Pax7^+^ or MyoD^+^ cells between control and IRE1α knockdown cultures **(Fig. S1C-F)**. Western blot analysis also showed that knockdown of IRE1α does not affect the levels of Pax7 or MyoD protein in cultured myoblasts (**Fig. S1G**). We have previously reported that c-Jun N-terminal kinase (JNK) phosphorylates c-Jun transcription factor, which in turn induces the gene expression of *Pax7* in cultured myoblasts (28). Interestingly, in conditions of ER stress, IRE1α interacts with TRAF2 to phosphorylate ASK1 which in turn phosphorylates and activates JNK1/2 (16). Consistent with the levels of Pax7, there was no difference in the levels of phosphorylated or total JNK between control and IRE1α knockdown cultures **(Fig. S1G)**. Furthermore, knockdown of IRE1α did not affect the proliferation of cultured myoblasts measured by EdU incorporation assay **(Fig. S1H, I)**. Collectively, these results suggest that satellite cell-specific deletion of IRE1α inhibits myofiber regeneration without affecting the proliferation or differentiation of satellite cells.

### Targeted deletion of IRE1α inhibits myoblast fusion

We next investigated whether targeted ablation of IRE1α affects fusion of muscle precursor cells with injured myofibers. TA muscle of *Ern1^fl/fl^* and *Ern1^scKO^* mice was injured using intramuscular injection of 1.2 % BaCl_2_ solution. After 72 h, the mice were given single intraperitoneal injection of EdU. Finally, TA muscle was collected on day 14 post-injury, followed by analysis of EdU positive nuclei. Results showed that the number of EdU^+^ nuclei per myofiber and the proportion of myofibers containing two or more centrally located EdU^+^ nuclei were significantly reduced in injured TA muscle of *Ern1^scKO^* mice compared to corresponding *Ern1^fl/fl^* mice **(Fig. 4A-C)** suggesting fusion defects in *Ern1^scKO^* mice. Next, we investigated whether IRE1α affects fusion of cultured myoblasts. Primary myoblasts prepared from WT mice were transfected with control or IRE1α siRNA. The cells were then incubated in DM for 24 or 48 h, followed by immunostaining for myosin heavy chain (MyHC) protein and DAPI staining. Results showed that knockdown of IRE1α significantly reduced the number of myotubes (containing two or more nuclei) at 24 h after addition of DM **(Fig. 4D, E)**. At 48 h of addition of DM, we found that the proportion of myotubes containing 2-4 nuclei was significantly increased whereas proportion of myotubes containing 5-10 nuclei or more than 10 nuclei was significantly reduced in IRE1α knockdown cultures compared to control cultures (**Fig. 4D, F**). Indeed, the number of MyHC^+^ mononucleated cells was also significantly increased in IRE1α knockdown cultures compared to corresponding control cultures at 48 h of incubation in DM (**Fig. 4G**). Moreover, average myotube diameter was also significantly reduced in IRE1α knockdown cultures compared to control cultures **(Fig. 4H)**. Western blot analysis showed that while the levels of MyoD and MyHC were comparable, the levels of IRE1α protein were drastically reduced in myoblast cultures transfected with IRE1α siRNA compared to control cultures at 24 h of addition of DM **(Fig. S2A)**.

**FIGURE 4.**
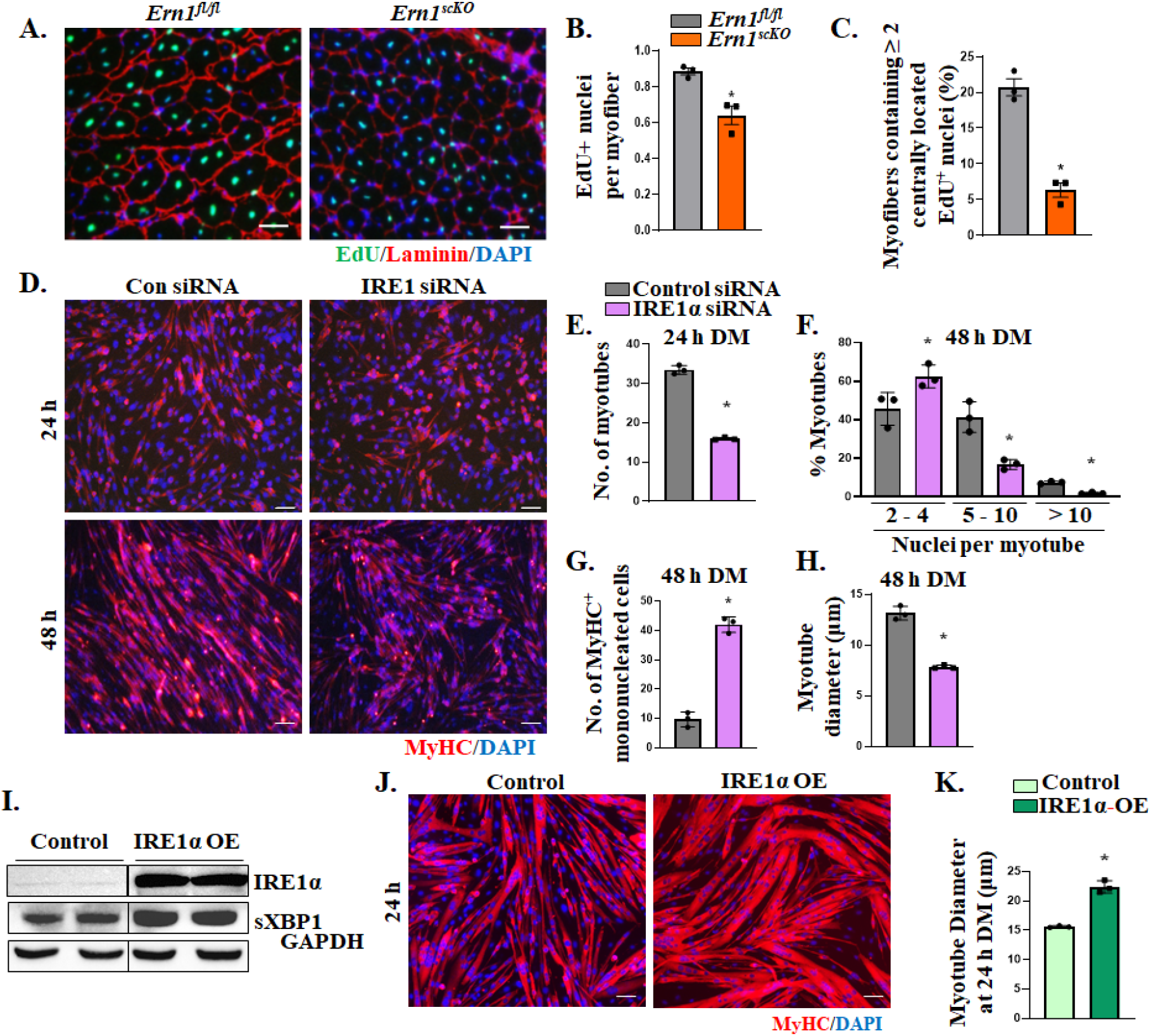
IRE1α is required for myoblast fusion both in vivo and in vitro. TA muscle of *Ern1^fl/fl^* and *Ern1^scKO^* mice was injured by intramuscular injection of 1.2% BaCl_2_ solution. An intraperitoneal injection of EdU was given after 3 days of injury and the muscle was collected on day 14 post-injury. **(A)** Transverse sections of injured TA muscle were processed for detection of EdU and immunostained with laminin to mark myofibers boundaries. DAPI was used to identify nuclei. Representative photomicrographs for stained sections of injured TA of *Ern1^fl/fl^* and *Ern1^scKO^* mice. Scale bar: 50 µm. Quantification of **(B)** EdU^+^ nuclei per myofiber and **(C)** percentage of myofibers containing 2 or more centrally located EdU^+^ nuclei. **(D)** Primary myoblasts isolated from WT mice were transfected with control or IRE1α siRNA for 24 h followed by incubation in differentiation medium (DM). Cells were fixed at 24 or 48 h after addition of DM and analyzed for myotube formation by performing immunostaining for MyHC. Nuclei were stained using DAPI. Scale bar: 50 µm. Quantitative analysis of **(E)** number of myotubes at 24 h, **(F)** percentage of myotubes containing 2-4, 5-10 and more than 10 nuclei at 48 h, **(G)** number of MyHC^+^ mononucleated cells at 48 h, and **(H)** average myotube diameter in control and IRE1α siRNA transfected cultures at 48 h after incubation in DM. **(I)** Primary myoblasts were stably transduced with control or IREα overexpressing (OE) retrovirus. Representative immunoblots showing protein levels of IRE1α and sXBP1, and an unrelated protein GAPDH in control and IRE1α OE cultures. Black lines on the immunoblots indicate that intervening lanes have been spliced out. **(J)** Control and IRE1α OE cells were incubated in DM for 24 h followed by immunostaining for MyHC protein and DAPI staining to analyze myotube formation. Scale bar: 50 µm. **(K)** Average diameter of MyHC^+^ cells. n=3 per group. Data are presented as mean ± SEM. *p ≤ 0.05, values significantly different from corresponding muscle of *Ern1^fl/fl^*mice or control cultures analyzed by unpaired Student *t* test.

We next investigated the effects of forced activation of IRE1α in cultured myoblasts. Previous studies have demonstrated that overexpression of IRE1α is sufficient for its activation in cultured mammalian cells in the absence of ER stress (15, 29). Primary myoblasts were transduced with retrovirus expressing a cDNA encoding IRE1α or enhanced green fluorescence protein (EGFP, as control). By performing western blot, we first confirmed that the levels of IRE1α protein are increased in cultures transduced with IRE1α cDNA encoding retrovirus compared to those transduced with EGFP retrovirus. There was also a considerable increase in the levels of sXBP1 protein in IRE1α overexpressing (OE) cultures **(Fig. 4I)**. Next, control and IRE1α OE cultures were incubated in DM for 24 h followed by immunostaining for MyHC protein. Results showed that overexpression of IRE1α improved formation of myotubes with an increased diameter **(Fig. 4J, 4K)**.

Recently, a pharmacological compound named IXA4 was identified as an efficient and selective activator of IRE1α-sXBP1 axis (30). We investigated whether pharmacological activation of IRE1α using IXA4 compound increases fusion of cultured myoblasts. We first determined the concentration of IXA4 that is required for augmenting the levels of sXBP1 protein in cultured myoblasts (**Fig. S2B**). Finally, primary myoblasts were treated with IXA4 and incubated in DM for 48 h followed by immunostaining for MyHC protein. Intriguingly, the average myotube diameter was significantly increased in cultures treated with IXA4 compared to corresponding controls treated with vehicle alone **(Fig. S2C, D)**. In another experiment, we also investigated whether IRE1α-mediated signaling is needed in one or both fusion partners during myogenesis. For this experiment, we generated lentiviral particles expressing either a scrambled shRNA or IRE1α shRNA along with mCherry protein. In addition, we used lentiviral particles expressing GFP protein alone. Primary myoblasts were stably transduced with lentiviral particles expressing GFP protein alone (control myoblasts) or scrambled or IRE1α shRNA along with mCherry. Finally, equal number of control and scrambled shRNA or IRE1α shRNA-expressing myoblasts were mixed and incubated in DM for 48 h. Results showed that knockdown of IRE1α in myoblasts considerably reduced their fusion with control myoblasts (**Fig. S2E**), suggesting that IRE1α is required in both fusion partners for efficient myotube formation.

Non-canonical NF-κB and canonical Wnt signaling play important roles in myoblast fusion during myogenesis (31–34). To understand the signaling mechanisms by which IRE1α promotes myoblast fusion, we investigated the effects of knockdown of IRE1α on the markers of canonical and non-canonical NF-κB and Wnt signaling at 48 h of incubation in DM. However, no difference was observed in the levels of MyHC (marker of muscle differentiation), p-p65 and p65 (markers of canonical NF-κB), p100 and p52 (markers of non-canonical NF-κB) and p-GSK3β (a marker of Wnt signaling) protein in control and IRE1α knockdown cultures. Furthermore, there was no significant difference in mRNA levels of Wnt ligands (Wnt4, Wnt5a, Wnt3a, and Wnt11), Wnt receptors (Fzd2, Fzd4, and Fzd6) and target gene *Axin2* in control and IRE1α knockdown cultures **(Fig. S2F, G)**. These results suggest that IRE1α promotes myoblast fusion without affecting the activation of NF-κB and Wnt signaling during myogenesis.

### Transcriptome analysis corroborates the role of IRE1α in myoblast fusion

By performing RNA sequencing (RNA-Seq), we next studied global gene expression changes in control and IRE1α knockdown myoblast cultures at 24 h after the addition of DM. Differentially Expressed Genes (DEGs) were classified based on at least 1.5-times fold change from the basal levels of control cells coupled with the criteria for significance of p<0.05. Analysis of DEGs revealed that 521 genes were downregulated whereas 555 genes were upregulated in IRE1α knockdown cultures compared with controls **(Fig. 5A)**. Pathway analysis using Metascape Gene Annotation and Analysis Tool showed the involvement of downregulated gene sets in the processes, such as regulation of protein exit from ER, response to topologically incorrect proteins, and vesicle localization. By contrast, upregulated gene sets were associated with the regulation of nuclease activity, muscle system process, negative regulation of response to external stimulus, and regulation of vasculature development and angiogenesis **(Fig. 5B)**. Downregulation of protein-folding related genes comply with the established role of IRE1α in the resolution of ER stress (35). ER stress-mediated activation of IRE1α exerts its downstream effects through three identified mechanisms: JNK-mediated signaling, XBP1 mRNA splicing, and RIDD pathway (13, 29, 36, 37). Our preceding results showed that knockdown of IRE1α does not affect the activation of JNK in cultured myoblasts **(Fig. S1G)**. In addition, we found no significant changes in the transcript levels of target mRNAs of RIDD pathway, including *Hgsnat*, *Pdqfrb*, *Scara3*, and *Sparc* **(Fig. S3A).** Recent studies have identified a few mRNA in skeletal muscle and other cell types subjected to RIDD pathway (38). Our RNA-Seq analysis showed no significant alterations in selected putative substrates of RIDD pathway **(Fig. S3A),** suggesting that IRE1α does not regulate RIDD pathway during myogenic differentiation. Therefore, we focused our further investigation to determine whether knockdown of IRE1α impairs myoblast fusion through sXBP1-mediated transcriptional regulation. It is known that sXBP1 regulates the gene expression of several molecules which are involved in multiple pathways, including but not limited to protein folding, protein entry into the ER, and ER-associated degradation (ERAD) pathway (39, 40). Heatmaps generated using z-scores based on transcript per million (TPM) values showed diminished transcript levels of multiple molecules associated with the ERAD pathway **(Fig. 5C)**. We next investigated whether IRE1α regulates the gene expression of ERAD molecules in myoblasts through XBP1 transcription factor. For this experiment, primary myoblasts were transfected with control or XBP1 siRNA followed by incubation in DM for 24h and performing RNA-Seq analysis. Remarkably, the mRNA levels of majority of ERAD-related molecules downregulated in IRE1α knockdown cultures were reduced in XBP1 knockdown myoblast cultures **(Fig. S3B)**.

**FIGURE 5.**
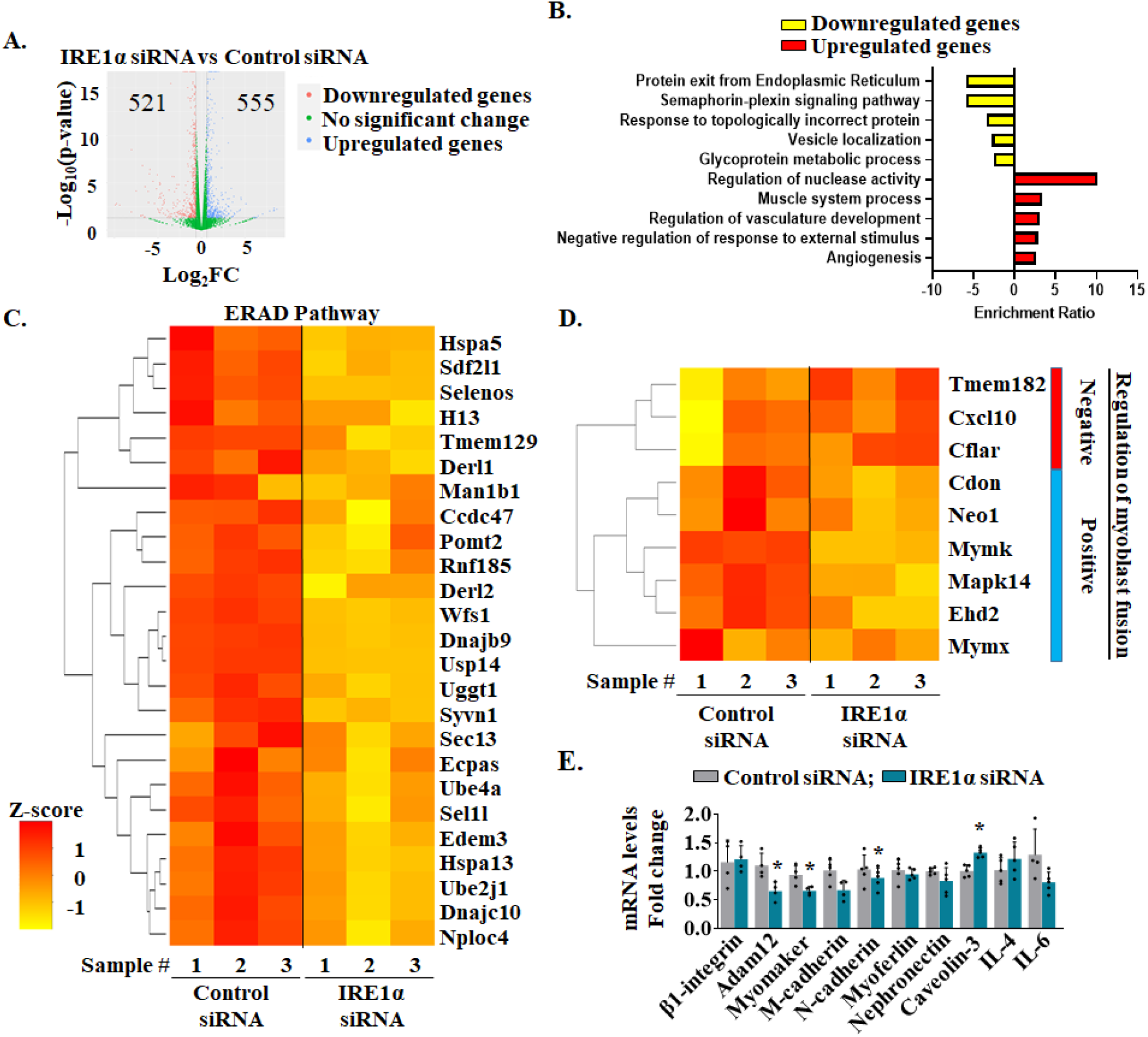
Global transcriptomic changes in IRE1α deficient myoblast cultures. Mouse myoblasts were treated with control or IRE1α siRNA followed by incubation in DM for 24 h. RNA was isolated from these cultures and used for RNA-Sequencing analysis. Differentially expressed genes (DEGs) were sorted by using the criteria of Log2FC ≥ |0.5| and significance of p ≤ 0.05. **(A)** Volcano plot showing the distribution of upregulated, downregulated and significantly unaltered genes. **(B)** Upregulated and downregulated gene sets were subjected to pathway analysis using Metascape Gene Annotation and Analysis tool. Enriched Gene Ontology (GO) Biological processes and Reactome pathways associated with the mapped genes are presented here. **(C)** Heatmap representing z-scores (based on transcript per million reads) of genes associated with the ERAD pathway compared between control- and IRE1α-siRNA groups. **(D)** Heatmap representing selected genes involved in the regulation of myoblast fusion. **(E)** Transcript levels of pro-fusion molecules control and IRE1α knockdown cultures at 24 h of incubation in DM assayed by performing qRT-PCR. N=3 per group. Data are presented as mean ± SEM. *p ≤ 0.05, values significantly different from control cultures analyzed by unpaired Student *t* test.

We next investigated whether knockdown of IRE1α or XBP1 affects gene expression of molecules that mediate myoblast fusion. We found that siRNA-mediated knockdown of IRE1α or XBP1 resulted in deregulation of both positive and negative regulators of myoblast fusion (**Fig. 5D, S3C**). Interestingly, mRNA levels of Myomaker (*Mymk*), a critical regulator of myoblast fusion, were significantly reduced in IRE1α or XBP1 knockdown cultures compared to corresponding controls **(Fig. 5D, Fig. S3C)**. In a separate experiment, we also measured relative mRNA levels of a few other profusion molecules in control or IRE1α knockdown myoblast cultures at 24 h of incubation in DM. Results showed that mRNA levels of Adam12, Myomaker, and N-cadherin were significantly reduced, whereas mRNA levels of Caveolin-3 were increased in IRE1α knockdown cultures compared to controls **(Fig. 5E)**. We also investigated the effect of overexpression of IRE1α on the transcript levels of various profusion molecules. Interestingly, the mRNA levels of Myomaker, myomerger, M-cadherin, Caveolin-3, β1D-integrin, and nephronectin were significantly increased in IRE1α overexpressing myoblasts compared to controls at 24 h of addition of DM (**Fig. S4A**). Collectively, these results suggest that IRE1α-mediated signaling promotes myoblast fusion through augmenting the gene expression of specific profusion molecules, including Myomaker.

### IRE1α augments levels of sXBP1 and Myomaker during myogenesis

We next investigated whether IRE1α promotes myoblast fusion through augmenting the levels of transcriptionally active sXBP1 protein. Primary myoblasts prepared from WT mice were transfected with control or XBP1 siRNA followed by incubation in DM for 24 or 48 h. The myotube formation was evaluated by immunostaining for MyHC protein **(Fig. 6A)**. Interestingly, myotube formation was significantly reduced in myoblast cultures transfected with XBP1 siRNA at 24 h after incubation in DM **(Fig. 6A, B)**. At 48 h, the number of myotubes with 2-4 nuclei was significantly increased whereas the number of myotubes containing 5-10 or more than 10 nuclei was significantly reduced in cultures transfected with XBP1 siRNA compared to control cultures (**Fig. 6C**). Moreover, the number of MyHC^+^ mononucleated cells was significantly increased in XBP1 siRNA transfected cultures **(Fig. 6D)**. In addition, our analysis showed that there was a significant reduction in average myotube diameter in XBP1 knockdown cultures compared to control cultures at 48 h of addition of DM **(Fig. 6A, E)**. Western blot analysis showed that the protein levels of MyHC were comparable between control and XBP1 knockdown cultures (**Fig. S4B**), suggesting that siRNA-mediated knockdown of XBP1 inhibits myotube formation without affecting myogenic differentiation.

**FIGURE 6.**
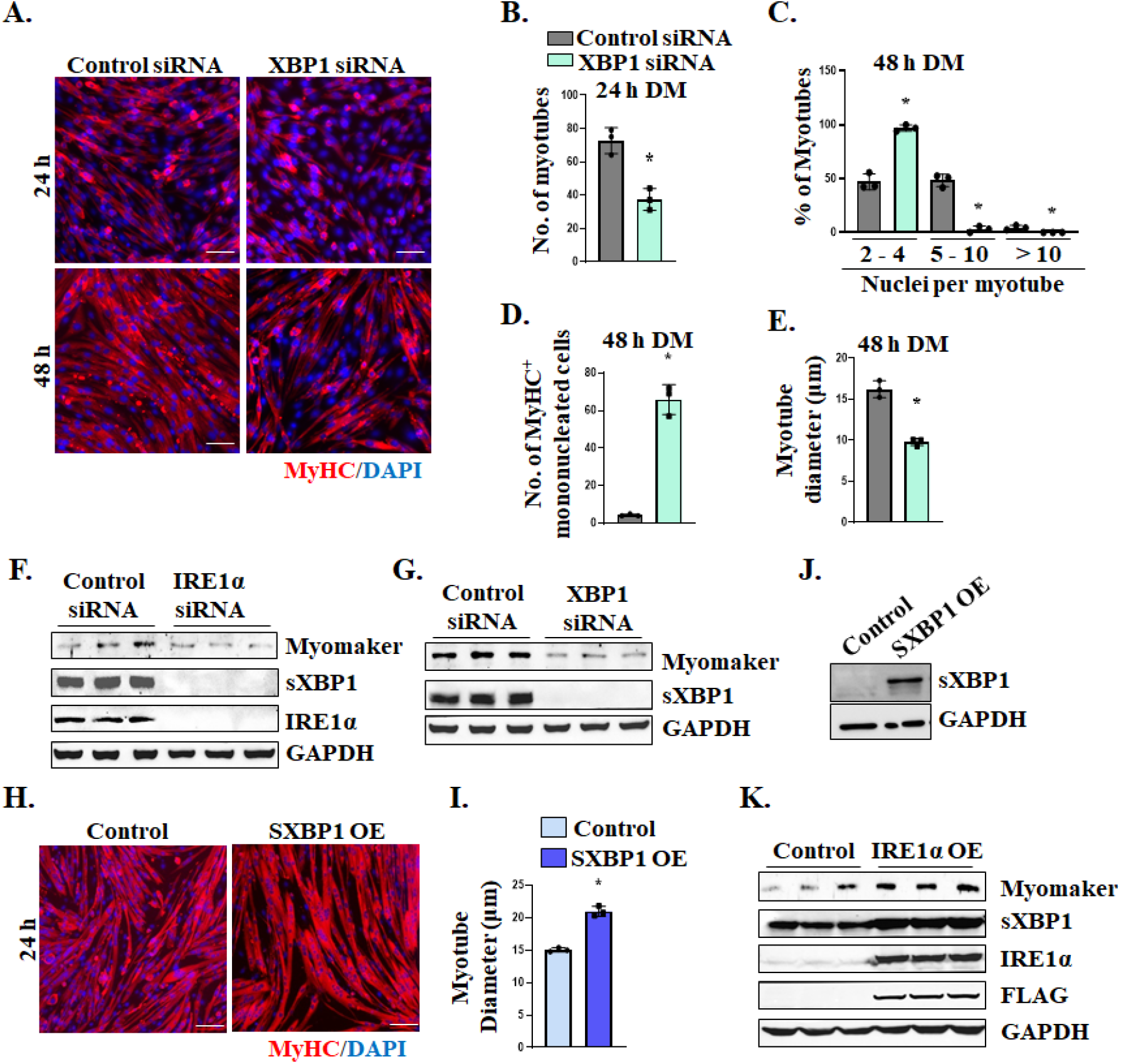
IRE1α regulates myoblast fusion through XBP1. Primary myoblasts were transfected with control or XBP1 siRNA for 24h. The cultures were then incubated in DM for 24 or 48 h followed by immunostaining for MyHC protein. DAPI was used to stain nuclei. Representative photomicrographs are presented here. Scale bar: 50µm. Quantitative analysis of **(B)** number of myotubes at 24 h, **(C)** percentage of myotubes containing 2-4, 5-10 or more than 10 nuclei at 48 h, **(D)** number of MyHC^+^ mononucleated cells at 48 h, and **(E)** average myotube diameter at 48 h of DM in control and XBP1 knockdown cultures. n=3 per group. Data are presented as mean ± SEM. *p ≤ 0.05, values significantly different from control cultures analyzed by unpaired Student *t* test. **(F)** Representative immunoblots presented here show protein levels of Myomaker, sXBP1, IRE1α, and unrelated protein GAPDH in cultures transfected with control or IRE1α siRNA and incubated in DM for 24 h. **(G)** Representative immunoblots presented here show the levels of Myomaker, sXBP1, and GAPDH in cultures transfected with control or XBP1 siRNA at 24 h of incubation in DM. **(H)** Primary myoblasts were transduced with control or sXBP1 cDNA expressing retrovirus for 48h. The cells were then incubated in DM for 24 h followed by immunostaining for MyHC protein. DAPI was used to stain nuclei. Representative photomicrographs presented here show formation of thicker myotubes in sXBP1 OE cultures. Scale bar: 50 µm. **(I)** Quantification of average myotube diameter in control and sXBP1 OE cultures. **(J)** Western blot showing protein levels of sXBP1 and unrelated protein GAPDH in control and sXBP1 OE cultures. **(K)** Primary myoblasts were transduced with control or IRE1α OE retroviral particles followed by incubation in DM for 24 h. Representative immunoblots presented here show the levels of Myomaker, sXBP1, IRE1α, FLAG, and unrelated protein GAPDH.

Transmembrane protein Myomaker plays an essential role in myoblast fusion (7, 10). Our RNA-Seq analysis revealed that mRNA levels of Myomaker are reduced upon knockdown of IRE1α or XBP1 in cultured myoblasts. We next investigated whether IRE1α increases the expression of Myomaker through augmenting levels of sXBP1 protein. Results showed that siRNA-mediated knockdown of IRE1α reduced the levels of both sXBP1 and Myomaker protein in cultured myoblasts incubated in DM for 24 h (**Fig. 6F**). Similarly, siRNA-mediated knockdown of XBP1 also reduced the amount of Myomaker protein in cultured myoblast at 24 h of incubation in DM (**Fig. 6G**). We next investigated the effect of overexpression of sXBP1 on myotube formation. Interestingly, retroviral-mediated overexpression of sXBP1 improved myotube formation at 24 h of incubation in DM (**Fig. 6H-J**). In another experiment, we investigated how overexpression of IRE1α affects the levels of sXBP1 and Myomaker protein in cultured myoblasts. Results showed that the levels of both Myomaker and sXBP1 protein were increased in IRE1α overexpressing cultured myoblasts (**Fig. 6K**). Collectively, these results suggest that the activation of IRE1α induces myoblast fusion through enhancing the levels of sXBP1 protein, which induces the gene expression of Myomaker.

### Transcription factor sXBP1 binds to the promoter region of *Mymk* gene

The sXBP1 protein binds to the promoter regions of multiple genes to regulate their transcription. We first determined whether the promoter region of *Mymk* gene contains consensus core DNA sequence for sXBP1 transcription factor. Specifically, the UCSC genome browser was used to locate and identify conserved binding domains of sXBP1 in the regulatory region of *Mymk* gene in the Mouse GRCm39/mm39 genome database. The ReMap ChIP-Seq function is an integrative atlas of multiple ChIP-Seq analyses deposited on Gene Expression Omnibus database (41–44). Interestingly, we observed a predicted sXBP1 binding site in the proximity of *Mymk* gene, which is also described by Liu *et al* (45) for ER transporter protein genes, such as *Sec23b*, *Sec61a1*, and *Sec61b*. To identify these genome sequences containing the core-binding domain of sXBP1, the GenBank sequence of the two predicted isoforms of *Mymk* gene, flanking the region 26951648-26962173 on chromosome 2 (GRCm39), were manually searched for the “ACGT” core binding sXBP1 domain **(Fig. 7A)**. We identified two sites, named Binding-Site1 and Binding-Site2 localized after exon1 of *Mymk* predicted isoform 1 or ∼4000 bp for Site1 and ∼2200 bp for Site2 upstream of *Mymk* predicted isoform 2 **(Fig. 7B)**. To determine whether sXBP1 binds to these sequences during myogenesis, we performed chromatin immunoprecipitation (ChIP) assay followed by semi-quantitative PCR and quantitative real time-PCR assays using primer sets designed to amplify 100-150 bp genome sequence flanking predicted DNA binding Site1 or Site2 in *Mymk* gene. Results showed increased enrichment of sXBP1 to both predicted sites in *Mymk* gene **(Fig. 7C, D)**. The ChIP experiment was confirmed using standard PCRs for positive control (Rpl30 for Histone H3 ChIP antibody) and known sXBP1 target sequence in promoter regions of Hspa5 and Dnajb9 genes **(Fig. 7E)**. Altogether, these results suggest that sXBP1 protein augments gene expression of Myomaker through directly binding to its promoter.

**FIGURE 7.**
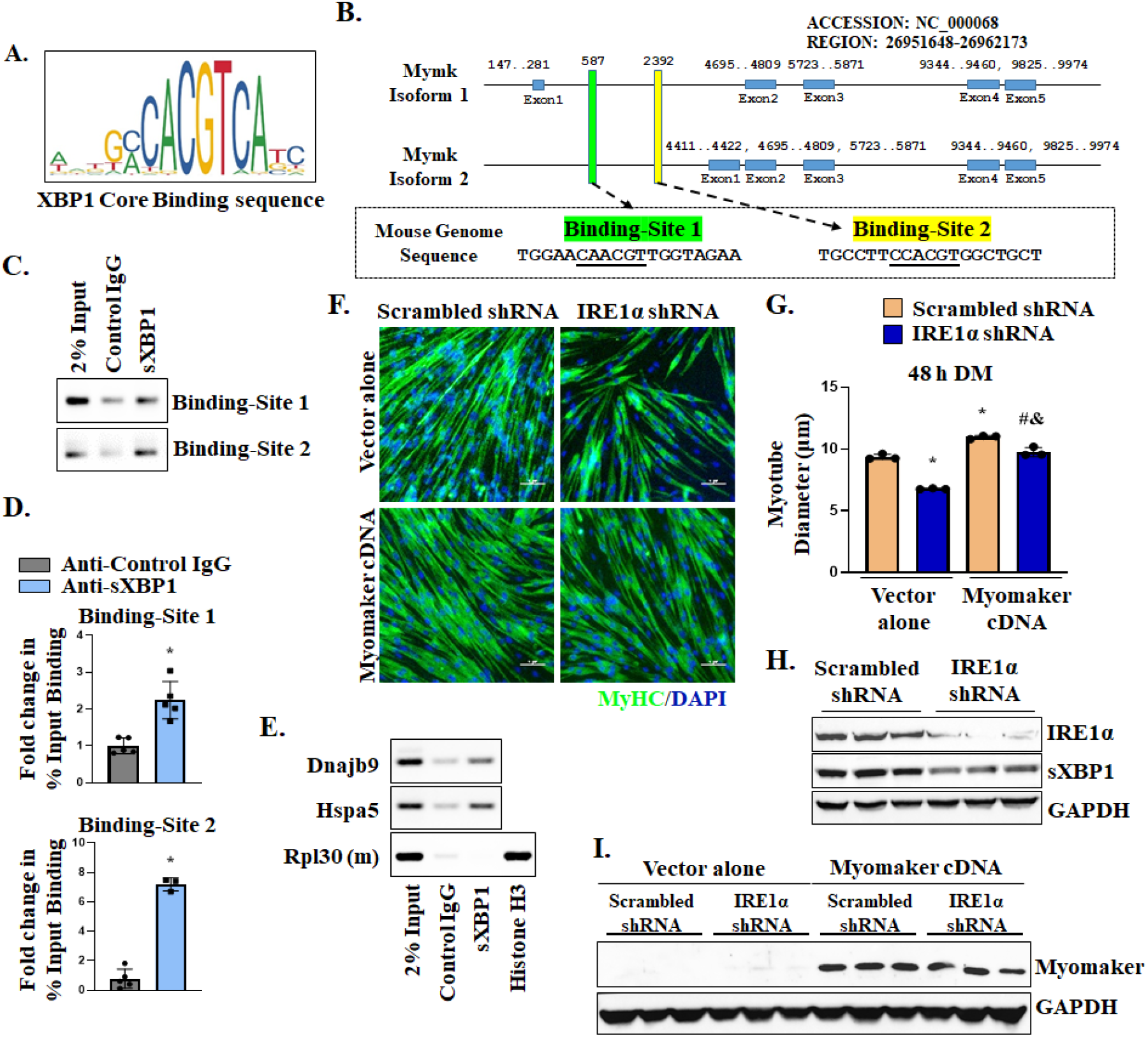
sXBP1 regulates the gene expression of Myomaker during myoblast fusion. **(A)** Symbolic representation of core binding sequence of sXBP1 transcription factor (downloaded from JASPER website). **(B)** Schematic representation of *Mymk* gene sequence in the mouse genome including Mymk-Site1 and Mymk-Site2, which contain sXBP1 core binding sequence. **(C)** Primary myoblasts were processed for ChIP assay for the binding of sXBP1 at its putative consensus DNA sequence in *Mymk* gene. Agarose gel images of semi-quantitative RT-PCR show levels of enrichment of *sXBP1* at indicated sXBP1 sites in mouse *Mymk* gene. **(D)** ChIP assay followed by qPCR analysis depicting percentage of input enrichment of sXBP1 at putative Site 1 and Site 2 in Mymk gene in cultured myoblasts. n=3-5 per group. Data are presented as mean ± SEM. *p ≤ 0.05; values significantly different from control IgG analyzed by unpaired Student *t* test. **(E)** ChIP assay results were validated using standard PCR reactions for positive control Rpl30 against Histone H3 antibody and known sXBP1 targets Dnajb9 and Hspa5. **(F)** Myoblasts stably expressing scrambled or IRE1α shRNA were transfected with empty vector or plasmid containing Mymk cDNA. After 24 h, the cultures were incubated in DM for 48 h and analyzed for myotube formation. Representative photomicrographs of the cultures after immunostaining for MyHC protein. DAPI was used to counterstain nuclei. Scale bar: 50µm. **(G)** Quantification of myotube diameter in control and Myomaker overexpressing cultures. n=3 per group. Data are presented as mean ± SEM. *p ≤ 0.05, values significantly different from corresponding control cultures analyzed by unpaired Student *t* test. **(H)** Western blot analysis demonstrating knockdown of IRE1α and reduced levels of sXBP1 in cultures expressing IRE1α shRNA. **(I)** Western blot confirming elevated levels of Myomaker protein in cultures transfected with *Mymk* cDNA.

To understand whether IRE1α-XBP1 signaling induces myoblast fusion through augmenting the gene expression of Myomaker, we investigated whether forced expression of Myomaker can rescue fusion defects in IRE1α knockdown cultures. Interestingly, reintroduction of *Mymk* significantly improved myotube diameter in IRE1α knockdown cultures **(Fig. 7F-I)**. Collectively, these results suggest that the IRE1α-XBP1 signaling promotes myoblast fusion through augmenting the gene expression of Myomaker.

### IRE1α regulates myoblast fusion during overload-induced muscle growth

In addition to muscle regeneration, myoblast fusion is also essential for myofiber hypertrophy and skeletal muscle growth in response to functional overload (34, 46). To confirm the physiological significance of IRE1α/XBP1 signaling in myoblast fusion in adult mice, we investigated whether IRE1α/XBP1 signaling also promotes myoblast fusion during overload-induced muscle hypertrophy. We used a bilateral synergistic ablation (SA) model that has been consistently used to induce hypertrophy of the plantaris muscle of adult mice (34, 47). After 2 days of sham or SA surgery, the mice were given an intraperitoneal injection of EdU and plantaris muscle was collected on day 14 of SA surgery. There was a significant increase in the wet weight of plantaris muscle in both *Ern1^fl/fl^* and *Ern1^scKO^* mice at 14d of performing SA surgery. However, gain in wet weight of plantaris muscle in response to functional overload normalized by body weight was significantly reduced in *Ern1^scKO^* mice compared with *Ern1^fl/fl^* mice (**Fig. 8A**). Next, we generated transverse sections of plantaris muscle and performed H&E staining followed by analysis of myofiber CSA. Results showed that average myofiber CSA of 14d-overloaded plantaris muscle was significantly reduced in *Ern1^scKO^* mice compared to *Ern1^fl/fl^* mice **(Fig. 8B, C)**. To determine whether reduction in myofiber CSA in plantaris muscle of *Ern1^scKO^* mice in response to functional overload is attributed to deficit in myoblast fusion, we analysed plantaris muscle sections for EdU incorporation in myofibers (**Fig. 8D**). Interestingly, the percentage of myofibers containing EdU^+^ nuclei in plantaris muscle was significantly reduced in Ern1^scKO^ mice compared to *Ern1^fl/fl^* mice (**Fig. 8E**). These results further suggest that IRE1α promotes skeletal muscle growth in adult mice through augmenting myoblast fusion.

**FIGURE 8.**
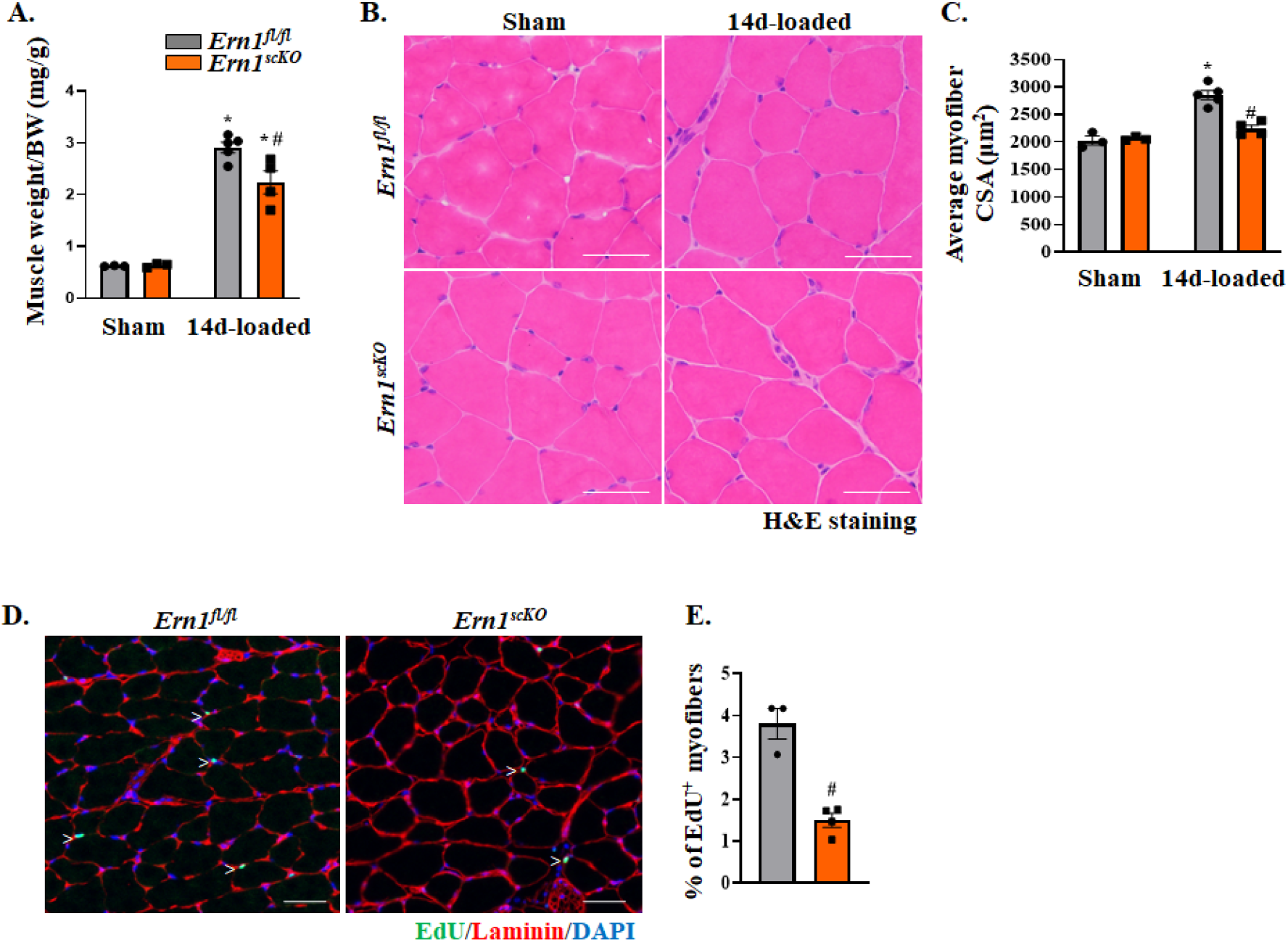
IRE1α promotes myoblast fusion during overload-induced muscle hypertrophy. *Ern1^fl/fl^* and *Ern1^scKO^* mice were subjected to sham or synergistic ablation (SA) surgery to put functional overload on plantaris muscle. After 3d, the mice were given a single intraperitoneal injection of EdU. At 14d of SA surgery, the mice were euthanized and plantaris muscle was isolated and analyzed. **(A)** Quantification of wet weight of plantaris muscle normalized to body weight of *Ern1^fl/fl^* and *Ern1^scKO^* mice. **(B)** Representative photomicrographs of H&E stained transverse sections of sham or 14d-loaded plantaris muscle of *Ern1^fl/fl^* and *Ern1^scKO^* mice. Scale bar: 50 µm. **(C)** Quantitative analysis of average myofibers CSA of sham and 14d-loaded plantaris muscle of Ern1^fl/fl^ and Ern1^scKO^ mice. **(D)** Representative images 14d-loaded plantaris muscle of *Ern1^fl/fl^* and *Ern1^scKO^* mice after staining for EdU and laminin protein. DAPI was used to counterstain nuclei. Scale bar: 50 µm. **(E)** Quantification of proportion of EdU^+^ myofibers in 14d-loaded plantaris muscle of *Ern1^fl/fl^* and *Ern1^scKO^* mice. n=3-4 per group. Data are presented as mean ± SEM and analyzed by unpaired Student *t* test or two-way ANOVA followed by Tukey’s multiple comparison test. *p ≤ 0.05; values significantly different from sham-operated of *Ern1^fl/fl^* mice. #p ≤ 0.05; values significantly different from 14d-overloaded plantaris muscle of *Ern1^fl/fl^* mice.

## DISCUSSION

Skeletal muscle regeneration is regulated by a complex network of signaling pathways. In addition to pro-myogenic signaling, accumulating evidence suggests that skeletal muscle regeneration involves the activation of UPR pathways (18, 19, 23). We recently reported that the IRE1α/XBP1 arms of the UPR is highly activated in skeletal muscle in response to injury and that myofiber-specific deletion of IRE1α or XBP1 attenuates muscle regeneration potentially through inhibiting the proliferation of satellite cells in a cell non-autonomous manner (23). However, the cell autonomous role and mechanisms of action of IRE1α in muscle progenitor cells remained unknown. In the present study, our results demonstrate that IRE1α promotes myoblast fusion through enhancing the levels of sXBP1 protein and augmenting the gene expression of Myomaker, a critical regulator of myoblast fusion.

Recent studies from our group and other have highlighted the role of UPR pathways in muscle progenitor cell function and myogenic differentiation. It has been found that the PERK/eIF2α arm of the UPR is essential for the survival and self-renewal of satellite cells in skeletal muscle of adult mice (18, 21, 22). Loss of PERK drastically reduces the pool of satellite cells in injured myofibers resulting in a significant decrease in various parameters of muscle regeneration, including the gene expression of myogenic regulator factors (MRFs), MyoD and Myogenin and the levels of eMyHC (21). In contrast, satellite cell-specific ablation of XBP1 in mice did not affect the expression of MRFs or eMyHC in regenerating skeletal muscle suggesting that XBP1 does not affect the myogenic function of satellite cells in adults. Since the markers of muscle differentiation were not affected, we did not perform extensive analysis of muscle regeneration phenotype in satellite cell-specific XBP1-knockout mice. However, our limited analysis showed a trend towards reduction in average myofiber CSA and number of myofibers containing two or more centrally located nuclei in TA muscle of satellite cell-specific XBP1 knockout mice at day 5 post-injury (21).

Proliferating satellite cells eventually differentiate into myoblasts/myocytes, which fuse with each other or to damaged myofibers to accomplish muscle repair. Our results using loss-of-function and gain-of-function approaches demonstrate that IRE1α-mediated signaling promotes myoblast fusion (**Fig. 4** and **Fig. S2**). Myogenic differentiation is tightly controlled by the sequential expression of various MRFs. Consistent with results in satellite cell-specific XBP1-knockout mice (21), there was no significant difference in the protein levels of MyoD and myogenin in regenerating skeletal muscle of *Ern1^fl/fl^* and *Ern1^scKO^* mice. Similarly, knockdown of IRE1α in cultured myoblasts did not affect the levels of MyoD or myogenin protein at different time points after induction of differentiation **(Fig. 3)** suggesting that the deficiency of IRE1α or XBP1 in satellite cells specifically inhibits myoblast fusion without having any effect on the myogenic differentiation program.

Myoblast fusion is also critical for overload-induced myofiber hypertrophy in adults (46, 48, 49). Our results demonstrate that IRE1α also promotes myoblast fusion during overload-induced muscle growth (**Fig. 8**). Whole genome RNA-Seq analysis of IRE1α knockdown myoblasts showed gene enrichment of negative regulators of myoblast fusion. In addition, transcript levels of multiple profusion molecules are reduced in IRE1α knockdown cultured myoblasts **(Fig. 5)**. Furthermore, we demonstrate that IRE1α promotes myoblast fusion through increasing the levels of sXBP1, which regulates the gene expression of profusion molecules. This is evidenced by the findings that knockdown of XBP1 in cultured myoblasts also inhibits their fusion and results in the dysregulation of various molecules involved in myoblast fusion. More importantly, forced expression of IRE1α or sXBP1 in cultured myoblasts is sufficient to induce expression of specific profusion molecules and facilitate myotube formation (**Fig. 6, Fig. S3**).

Myomaker, a membrane protein, is critical for myoblast fusion. A recent study demonstrated that promoter of Mymk contains E-Box motifs to which MyoD transcription factor binds to induce the gene expression of Myomaker for myotube formation (9). The results of the present study suggest the IRE1α-XBP1 signaling axis promotes myoblast fusion through transcriptional upregulation of *Mymk* gene **(Fig. 6)**. Transcriptionally active sXBP1 transcription factor regulates gene expression of target molecules by interacting with the “ACGT” core-binding domain within their promoter or regulatory region. Several target genes regulated by sXBP1 have been identified that play important roles in ER proteostasis (50, 51). We have identified two sXBP1 binding sites in mouse *Mymk* gene and validated enrichment of sXBP1 to these sites during myogenic differentiation. Furthermore, our loss-of-function and gain-of-function approaches demonstrate that IRE1α/XBP1 regulates gene expression of Myomaker and that overexpression of Myomaker is sufficient to rescue fusion defects in IRE1α knockdown myogenic cultures **(Fig. 7)**. Interestingly, our RNA-Seq analysis also showed that knockdown of IRE1α in cultured myoblasts leads to the dysregulation of expression of a few other positive and negative regulators of myoblast fusion (**Fig. 5D**). However, the mechanisms by which IRE1α regulates the levels of those molecules remain unknown. Future studies will investigate whether IRE1α-XBP1 signaling modulates the gene expression of other fusion-related molecules in a similar fashion as Myomaker during myogenesis.

Recently, He *et al.* showed that IRE1α signaling promotes muscle regeneration through RIDD-dependent decay of myostatin mRNA (52). In contrast, we found that IRE1α does not affect the mRNA levels of myostatin or other known RIDD substrates in regenerating skeletal muscle of mice (23). Instead, our RNA-Seq analysis of IRE1α knockdown cultures in this study showed reduction in the mRNA levels of myostatin (**Fig. S3A**). In addition, we did not find any significant difference in the early markers of myogenic differentiation, suggesting that IRE1α does not promote myoblast fusion through RIDD-dependent degradation of myostatin mRNA. Indeed, our results clearly show that IRE1α endonuclease activity causes the processing of XBP1 mRNA, which results in formation of transcriptionally active XBP1 transcription factor, leading to enhanced gene expression of profusion molecules.

While our experiments suggest a role of IRE1α/XBP1 signaling in myoblast fusion, it remains unknown whether this pathway is activated due to ER stress during myogenesis or through signaling crosstalk with other pathways. Interestingly, we found that in addition to profusion molecules, knockdown of IRE1α or XBP1 reduces mRNA levels of many molecules involved in the ER-associated degradation (ERAD) pathway (**Fig. 5C, Fig. S3**). It is possible that the activation of UPR pathways is essential for myogenesis and one of the critical functions of the IRE1/XBP1 arm of the UPR is to augment myoblast fusion (18, 19). Interestingly, our experiments showed that overexpression of IRE1α or sXBP1 in myoblasts does not induce fusion in growth conditions. Rather, myotube formation is improved when IRE1α or sXBP1-overexpressing myoblasts are incubated in differentiation medium suggesting that the activation of the UPR is a physiological response required for muscle formation and that distinct arm of the UPR regulates different aspects of myogenesis. While there is limited knowledge directly linking ERAD pathway with myoblast fusion, our results provide initial evidence that IRE1α-mediated regulation of ERAD-associated molecules could also facilitate myoblast fusion during muscle regeneration.

In summary, our results suggest that IRE1α/XBP1 signaling axis promotes myoblast fusion during regenerative myogenesis and overload-induced muscle growth in adult mice. While more investigation is needed to investigate mechanisms of action of IRE1α in myogenic cells, we provide initial evidence that augmenting the levels of IRE1α could improve muscle growth and repair in various muscle disorders.

## MATERIALS and METHODS

### Animals

Floxed *Ern1* (*Ern1^fl/fl^*) mice as described (53) were crossed with *Pax7-CreER* mice (Jax strain: B6;129-Pax7^tm2.1(cre/ERT2)Fan^/J) to generate satellite cell-specific *Ern1*-knockout (i.e. *Ern1^scKO^*) mice. Myofiber-specific Ern1-KO (*Ern1^mKO^*) mice have been described previously (23). All mice were in the C57BL6 background. Genotype of mice was determined by performing PCR from tail DNA. We used 9-16 weeks old mice for our experimentation. For Cre-mediated inducible deletion of *Ern1* in satellite cells, the mice were given intraperitoneal (i.p.) injection of tamoxifen (10 mg per Kg body weight) in corn oil for four consecutive days. *Ern1^fl/fl^* mice were also injected with tamoxifen following the same regimen and served as controls. All the animals were handled according to approved institutional animal care and use committee (IACUC) protocol (PROTO201900043) of the University of Houston. All surgeries were performed under anaesthesia, and every effort was made to minimize suffering. Details of all experimental procedures are described in the S1 Appendix Materials and Methods of Supplemental data file.

### Skeletal muscle injury and *in vivo* fusion assay

TA muscle of adult mice was injected 50 µl of 1.2 % BaCl_2_ (Sigma Chemical Co.) dissolved in saline to induce necrotic injury as described (28). The mice were euthanized at different time points after injury and TA muscle was isolated for biochemical and morphometric analysis. To study fusion of muscle progenitor cells *in vivo*, the mice were given an intraperitoneal injection of EdU (4 μg per gram body weight) at day 3 post BaCl_2_-medited injury of TA muscle. On day 11 after EdU injection, the mice were euthanized and TA muscle was isolated and transverse sections were made. The TA muscle sections were subsequently immunostained with anti-Laminin for marking boundaries of myofibers and processed for the detection of EdU^+^ nuclei. The EdU^+^ nuclei on muscle sections were detected as instructed in the Click-iT EdU Alexa Fluor 488 Imaging Kit (Invitrogen). DAPI stain was used to identify nuclei. Finally, images were captured and the number of intramyofiber EdU^+^ myonuclei per myofiber, percentage of 2 or more EdU^+^ centrally nucleated fibers, and percentage of EdU^+^ myonuclei/total DAPI^+^ nuclei were evaluated using NIH ImageJ software. Three to four different sections from mid-belly of each muscle were included for analysis.

### Synergistic ablation (SA) surgery

Bilateral SA surgery was performed in adult Ern1fl/fl and Ern1scKO mice following same procedure as described (34, 47). In brief, male mice were anesthetized using isoflurane and the soleus and ∼60 % of the gastrocnemius muscles were surgically removed while ensuring that the neural and vascular supply remained intact and undamaged for the remaining plantaris muscle. A sham surgery was performed for controls following exactly the same procedures except that gastrocnemius and soleus muscle were not excised. After 2d, the mice were given an intraperitoneal injection of EdU (4 μg/g body weight). At 14d of sham or SA surgery, the plantaris muscle was isolated and transverse muscle sections made were immunostained with anti-laminin and for detection of EdU and nuclei. The number of intrafiber EdU^+^ myonuclei/myofiber was quantified using NIH ImageJ software.

### Histology and morphometric analysis

Uninjured or injured TA muscle was isolated from mice and sectioned with a microtome cryostat. For the assessment of muscle morphology and quantification of myofiber cross-sectional area (CSA), 8-μm-thick transverse sections of TA muscle were stained with Hematoxylin and Eosin (H&E). The sections were examined under an Eclipse TE 2000-U microscope (Nikon, Tokyo, Japan). For quantitative analysis, average CSA of myofibers was analyzed in H&E-stained TA muscle sections using NIH ImageJ software. For each muscle, the distribution of myofiber CSA was calculated by analyzing approximately 250 myofibers.

### Immunohistochemistry

For immunohistochemistry studies, frozen TA or plantaris muscle sections were fixed in acetone or 4 % paraformaldehyde (PFA) in PBS, blocked in 2 % bovine serum albumin in PBS for 1 h followed by incubation with anti-Pax7 or anti-eMyHC and anti-laminin in blocking solution at 4 °C overnight under humidified conditions. The sections were washed briefly with PBS before incubation with goat anti-mouse Alexa Fluor 594 and goat anti-rabbit Alexa Fluor 488 secondary antibody for 1 h at room temperature and then washed three times for 15 min with PBS. Nuclei were counterstained with DAPI. The slides were mounted using fluorescence medium (Vector Laboratories) and visualized at room temperature on Nikon Eclipse TE 2000-U microscope (Nikon), a digital camera (Nikon Digital Sight DS-Fi1), and NIS Elements BR 3.00 software (Nikon). Image levels were equally adjusted using Photoshop CS6 software (Adobe).

### Myoblast isolation, culturing, and fusion assay

Primary myoblasts were prepared from the hind limbs of 8-week-old male or female mice following a protocol as described (34). To induce differentiation, the cells were incubated in differentiation medium (DM; 2 % horse serum in DMEM) for 24 or 48 h. The cultures were then fixed with 4 % PFA in PBS for 15 min at room temperature and permeabilized with 0.1 % Triton X-100 in PBS for 10 min. Cells were blocked with 2 % BSA in PBS and incubated with mouse anti-MyHC (clone MF20) overnight at 4 °C. The cells were then incubated with secondary antibody at room temperature for 1 h. Nuclei were counterstained with DAPI for 3 min. Stained cells were photographed and analyzed using a fluorescent inverted microscope (Nikon Eclipse TE 2000-U), a digital camera (Digital Sight DS-Fi1), and Elements BR 3.00 software (Nikon).

To measure fusion efficiency, more than one hundred MyHC^+^ myotubes containing 2 or more nuclei were counted. More than 5 field images were analyzed per experimental group. To measure average diameter of myotubes, 100–120 myotubes per group were included. For consistency, diameters were measured at the midpoint along with the length of the MyHC^+^ myotubes. Myotube diameter was measured using the NIH ImageJ software. Results presented are from 4–5 independent experiments. Image levels were equally adjusted using Photoshop CS6 software (Adobe).

### FACS

To detect p-IRE1α and IRE1α expression in cultured myoblasts, following the labeling with antibodies against CD45, CD31, Sca-1, Ter-119, and α7-integrin, the cells were fixed with 1 % PFA (Sigma-Aldrich) and permeabilized using 0.2 % Triton X-100 (Thermo Fisher Scientific). The cells were then incubated with anti-pIRE1α or anti-IRE1α and Alexa 488 secondary antibody (Invitrogen). FACS analysis was performed on a C6 Accuri cytometer (BD Biosciences) equipped with 3 lasers. The output data was processed, and plots were prepared using FCS Express 4 RUO software (De Novo Software).

### Viral vectors

pBABE-Puro (Addgene, Plasmid #1764), pBABE-Puro-EGFP (Addgene, Plasmid #128041), and Flag_HsIRE1a_pBabePuro (Addgene, Plasmid #54337) were purchased from Addgene (Watertown, MA). The pBX-8memC construct was provided by Dr. Douglas Millay of Cincinnati Children’s Hospital Medical Center, Cincinnati, OH. Mouse sXBP1 cDNA was isolated and ligated at BamHI and SalI sites in pBABE-Puro plasmid. The integrity of the cDNA was confirmed by performing DNA sequencing. For generation of retrovirus, about 5 x 10^6^ Platinum-E packaging cells (Cell Biolabs, Inc, San Diego, CA) were transfected with 5 μg of Flag_HsIRE1a_pBabePuro, pBABE-Puro-sXBP1, or pBABE-Puro-EGFP using FuGENE-HD (Promega, Madison, WI, USA). After 24 h of transfection, the medium was replaced with 10 % fetal bovine serum. Forty-eight hours after transfection, viral supernatants were collected, filtered through 0.45-micron filters, and then added to primary myoblasts in growth media containing 10 μg/ml polybrene. After two successive retroviral infections, cells were grown for 48 h and selected in the presence of 2 µg/ml puromycin.

The pLKO.1-mCherry-Puro plasmid was kindly provided by Dr. Renzhi Han of the Ohio State University. The target siRNA sequence was identified using BLOCK-iT™ RNAi Designer online software (Life Technologies). At least 2-3 siRNA sequence for IRE1α gene were tested for efficient knockdown of target mRNA. The shRNA oligonucleotides were synthesized to contain the sense strand of target sequences for mouse IRE1α (GCTAACGCCTACTCTGTATGT) short spacer (CTCGAG), and the reverse complement sequences followed by five thymidines as an RNA polymerase III transcriptional stop signal. Oligonucleotides were annealed and cloned into pLKO.1-mCherry-Puro with AgeI/EcoRI sites. The insertion of shRNA and cDNA sequence in the plasmids was confirmed by DNA sequencing. For generation of lentiviral particles, 5 x 10^6^ HEK293T cells were co-transfected with 5μg psPAX2 (Addgene, Plasmid # 12260), 5μg pMD2.G (Addgene, Plasmid # 12259) and 10μg of pLKO.1-mCherry-scrambled shRNA or pLKO.1-mCherry-IRE1α shRNA using FuGENE-HD reagent (Promega, Madison, WI, USA). After 24 h of transfection, the media was replaced with fresh media containing 10 % fetal bovine serum. Lentiviral particles were collected 48 h after transfection and filtered through 0.45-micron filters. Primary myoblasts were transduced with lentiviral particles containing scrambled or IRE1α-shRNA in growth medium containing 10 μg/ml polybrene. Cells were grown for 48 h and selected in the presence of 2 µg/ml puromycin.

### RNA isolation and qRT-PCR

RNA isolation and qRT-PCR were performed following a standard protocol as described (34). In brief, total RNA was extracted from uninjured and injured TA muscle of mice or cultured myoblasts using TRIzol reagent (Thermo Fisher Scientific) and RNeasy Mini Kit (Qiagen, Valencia, CA, USA) according to the manufacturers’ protocols. First-strand cDNA for PCR analyses was made with a commercially available kit (iScript cDNA Synthesis Kit, Bio-Rad Laboratories). The quantification of mRNA expression was performed using the SYBR Green dye (Bio-Rad SsoAdvanced - Universal SYBR Green Supermix) method on a sequence detection system (CFX384 Touch Real-Time PCR Detection System - Bio-Rad Laboratories). The sequence of the primers is described in **Supplemental Table S1**. Data normalization was accomplished with the endogenous control (β-actin), and the normalized values were subjected to a 2^-ΔΔCt^ formula to calculate the fold change between control and experimental groups.

### Chromatin Immunoprecipitation (ChIP) assay

Chromatin immunoprecipitation was performed using SimpleChIP enzymatic Chromatin IP kit (Cell Signaling Technology, Cat #9003) according to the manufacturers’ suggested protocol. Briefly, primary myoblasts were incubated in DM for 24 h followed by crosslinking, purification of nuclei, and chromatin shearing by sonication (8 times, 20s each). Sheared chromatin was incubated overnight with antibodies against sXBP1 (Cell Signaling Technology) or Histone H3 (Cell Signaling Technology) followed by incubation with Protein G magnetic beads. Normal Rabbit IgG (Cell Signaling Technology) was used as negative control for immunoprecipitation experiment. Magnetic beads were briefly washed, and chromatin DNA was eluted, reverse crosslinked, and purified. Purified DNA was analysed for enrichment of 100-150 bp sequences by quantitative real time-PCR (40 cycles) or semi-quantitative standard PCR using specific primers designed for binding sites in the regulatory regions of *Mymk*, *Hspa5*, *Dnajb9*, and mouse *Rpl30* (positive control) genes. Triplicates were used from two sets of experiments for standardization of qRT-PCR results. The fold change between negative control IgG and anti-sXBP1 groups was calculated using 2^-ΔΔCt^ formula. Primers used for PCR reactions are provided in **Supplemental Table S1**.

### Western Blot

TA muscle of mice or cultured primary myoblasts were washed with PBS and homogenized in lysis buffer (50 mM Tris-Cl (pH 8.0), 200 mM NaCl, 50 mM NaF, 1 mM dithiothreitol, 1 mM sodium orthovanadate, 0.3 % IGEPAL, and protease inhibitors). Approximately, 100 μg protein was resolved on each lane on 8-12% SDS-PAGE gel, transferred onto a nitrocellulose membrane, and probed using specific primary antibody (**Supplementary Table S2**). Bound antibodies were detected by secondary antibodies conjugated to horseradish peroxidase (Cell Signaling Technology). Signal detection was performed by an enhanced chemiluminescence detection reagent (Bio-Rad). Approximate molecular masses were determined by comparison with the migration of prestained protein standards (Bio-Rad). Uncropped gel images are presented in **Supplementary Fig. S5**.

### RNA sequencing and data analysis

Total RNA from control, IRE1α or XBP1 knockdown cultures was extracted using TRIzol reagent (Thermo Fisher Scientific) using the RNeasy Mini Kit (Qiagen, Valencia, CA, USA) according to the manufacturers’ protocols. The mRNA-seq library was prepared using poly (A)-tailed enriched mRNA at the UT Cancer Genomics Center using the KAPA mRNA HyperPrep Kit protocol (KK8581, Roche, Holding AG, Switzerland) and KAPA Unique Dual-indexed Adapter kit (KK8727, Roche). The Illumina NextSeq550 was used to produce 75 base paired end mRNA-seq data at an average read depth of ∼38M reads/sample. RNA-seq fastq data was processed using CLC Genomics Workbench 20 (Qiagen). Illumina sequencing adaptors were trimmed, and reads were aligned to the mouse reference genome Refseq GRCm39.105 from the Biomedical Genomics Analysis Plugin 20.0.1 (Qiagen). Normalization of RNA-seq data was performed using trimmed mean of M-values. Genes with fold change (FC) ≥ 1.5 (or log_2_FC ≥ 0.5) and FDR <0.05 were assigned as differentially expressed genes (DEGs) and represented in volcano plot using ggplot function in R software (v 4.2.2). Over-representation analysis was performed with a hypergeometric test using WebGestalt (v 0.4.3) (54). Gene Ontology (GO) Biological Processes associated with the upregulated and downregulated genes were identified with FDR cutoff value of 0.05. Network enrichment analysis was performed using Metascape Gene Annotation and Analysis tool (metascape.org) as described (55). Heatmaps were generated by using heatmap.2 function (56) using z-scores calculated based on transcripts per million (TPM) values. Average of absolute TPM values for control group are provided in **Supplemental Table S3**. TPM values were converted to log (TPM+1) to handle zero values. Genes involved in specific pathways were manually selected for heatmap expression plots. All the raw data files can be found on NCBI SRA repository using the accession code PRJNA1014488.

### Statistical analyses and experimental design

The sample size was calculated using power analysis methods for a priori determination based on the s.d. and the effect size was previously obtained using the experimental procedures employed in the study. For animal studies, we calculated the minimal sample size for each group as eight animals. Considering a likely drop-off effect of 10%, we set sample size of each group of six mice. For some experiments, three to four animals were sufficient to obtain statistically significant differences. Animals with same sex and same age were employed to minimize physiological variability and to reduce s.d. from mean. The exclusion criteria for animals were established in consultation with a veterinarian and experimental outcomes. In case of death, skin injury, ulceration, sickness, or weight loss of > 10%, the animal was excluded from analysis. Tissue samples were excluded in cases such as freeze artifacts on histological sections or failure in extraction of RNA or protein of suitable quality and quantity. We included animals from different breeding cages by random allocation to the different experimental groups. All animal experiments were blinded using number codes until the final data analyses were performed. Statistical tests were used as described in the Figure legends. Results are expressed as mean ± SEM. Statistical analyses used two-tailed Student’s *t*-test or 2-way ANOVA followed by Tukey’s multiple-comparison test to compare quantitative data populations with normal distribution and equal variance. A value of p < 0.05 was considered statistically significant unless otherwise specified.

## Data availability

All relevant data related to this manuscript are available from the authors upon reasonable request. All the raw data files for RNA-Seq experiment can be found on NCBI SRA repository using the accession code PRJNA1014488.

## Acknowledgements

This work was supported by National Institute of Health grants AR081487 and AR059810 to AK. We thank the technical support from the Cancer Prevention and Research Institute of Texas (CPRIT RP180734) for RNA-Seq experiment. We thanks Dr. Douglas Millay of Cincinnati Children’s Hospital Medical Center (Cincinnati, OH) for providing pBX-8memC construct. We also thank Dr. Pengpeng Bi of University of Georgia (Athens, GA) for providing myomaker antibody.

## Authors’ contribution

A.K. designed the work. A.S.J. and A.K. wrote the manuscript and all authors edited the manuscript. A.S.J, M.T.S., A.R., and T.E.K performed the experiments. T.I. provided reagents and edited the manuscript. M.B.C., M.W., Y.L. and P.H.G. helped with genomics experiments and bioinformatics analysis of RNA-Seq experiments.

## Conflict of Interest

The authors declare no competing interests.

## Notes

### Competing Interest Statement

The authors have declared no competing interest.

### Summary of Updates

We have added additional discussion in the results and discussion sections of the manuscript. Abstract has also been updated. Figure 3 is also revised. In addition, supplemental figure 3 is now removed. Data about role of Ern1 in muscle growth during neonatal stage is removed. This work will be included in a separate study.

## REFERENCES

1. Buckingham, M., Bajard, L., Chang, T., Daubas, P., Hadchouel, J., Meilhac, S., Montarras, D., Rocancourt, D., and Relaix, F. (2003) The formation of skeletal muscle: from somite to limb. J Anat 202, 59–68

2. Yin, H., Price, F., and Rudnicki, M. A. (2013) Satellite cells and the muscle stem cell niche. Physiol Rev 93, 23–67

3. Relaix, F., Bencze, M., Borok, M. J., Der Vartanian, A., Gattazzo, F., Mademtzoglou, D., Perez-Diaz, S., Prola, A., Reyes-Fernandez, P. C., and Rotini, A. (2021) Perspectives on skeletal muscle stem cells. Nature communications 12, 692

4. Abmayr, S. M., and Pavlath, G. K. (2012) Myoblast fusion: lessons from flies and mice. Development 139, 641–656

5. Kim, J. H., Jin, P., Duan, R., and Chen, E. H. (2015) Mechanisms of myoblast fusion during muscle development. Curr Opin Genet Dev 32, 162–170

6. Hochreiter-Hufford, A. E., Lee, C. S., Kinchen, J. M., Sokolowski, J. D., Arandjelovic, S., Call, J. A., Klibanov, A. L., Yan, Z., Mandell, J. W., and Ravichandran, K. S. (2013) Phosphatidylserine receptor BAI1 and apoptotic cells as new promoters of myoblast fusion. Nature 497, 263–267

7. Millay, D. P., O’Rourke, J. R., Sutherland, L. B., Bezprozvannaya, S., Shelton, J. M., Bassel-Duby, R., and Olson, E. N. (2013) Myomaker is a membrane activator of myoblast fusion and muscle formation. Nature 499, 301–305

8. Bi, P., Ramirez-Martinez, A., Li, H., Cannavino, J., McAnally, J. R., Shelton, J. M., Sanchez-Ortiz, E., Bassel-Duby, R., and Olson, E. N. (2017) Control of muscle formation by the fusogenic micropeptide myomixer. Science 356, 323–327

9. Zhang, H., Wen, J., Bigot, A., Chen, J., Shang, R., Mouly, V., and Bi, P. (2020) Human myotube formation is determined by MyoD-Myomixer/Myomaker axis. Sci Adv 6

10. Millay, D. P., Sutherland, L. B., Bassel-Duby, R., and Olson, E. N. (2014) Myomaker is essential for muscle regeneration. Genes & development 28, 1641–1646

11. Krauss, R. S. (2010) Regulation of promyogenic signal transduction by cell-cell contact and adhesion. Experimental cell research 316, 3042–3049

12. Adams, C. J., Kopp, M. C., Larburu, N., Nowak, P. R., and Ali, M. M. U. (2019) Structure and Molecular Mechanism of ER Stress Signaling by the Unfolded Protein Response Signal Activator IRE1. Front Mol Biosci 6, 11

13. Wang, M., and Kaufman, R. J. (2014) The impact of the endoplasmic reticulum protein-folding environment on cancer development. Nature Reviews Cancer 14, 581–597

14. Wu, J., and Kaufman, R. J. (2006) From acute ER stress to physiological roles of the unfolded protein response. Cell Death & Differentiation 13, 374–384

15. Acosta-Alvear, D., Karagoz, G. E., Frohlich, F., Li, H., Walther, T. C., and Walter, P. (2018) The unfolded protein response and endoplasmic reticulum protein targeting machineries converge on the stress sensor IRE1. Elife 7

16. Hetz, C. (2012) The unfolded protein response: controlling cell fate decisions under ER stress and beyond. Nat Rev Mol Cell Biol 13, 89–102

17. Hetz, C., Zhang, K., and Kaufman, R. J. (2020) Mechanisms, regulation and functions of the unfolded protein response. Nat Rev Mol Cell Biol 21, 421–438

18. Afroze, D., and Kumar, A. (2019) ER stress in skeletal muscle remodeling and myopathies. FEBS J 286, 379–398

19. Bohnert, K. R., McMillan, J. D., and Kumar, A. (2018) Emerging roles of ER stress and unfolded protein response pathways in skeletal muscle health and disease. J Cell Physiol 233, 67–78

20. Nakanishi, K., Dohmae, N., and Morishima, N. (2007) Endoplasmic reticulum stress increases myofiber formation in vitro. FASEB J 21, 2994–3003

21. Xiong, G., Hindi, S. M., Mann, A. K., Gallot, Y. S., Bohnert, K. R., Cavener, D. R., Whittemore, S. R., and Kumar, A. (2017) The PERK arm of the unfolded protein response regulates satellite cell-mediated skeletal muscle regeneration. Elife 6

22. Zismanov, V., Chichkov, V., Colangelo, V., Jamet, S., Wang, S., Syme, A., Koromilas, A. E., and Crist, C. (2016) Phosphorylation of eIF2alpha Is a Translational Control Mechanism Regulating Muscle Stem Cell Quiescence and Self-Renewal. Cell Stem Cell 18, 79–90

23. Roy, A., Tomaz da Silva, M., Bhat, R., Bohnert, K. R., Iwawaki, T., and Kumar, A. (2021) The IRE1/XBP1 signaling axis promotes skeletal muscle regeneration through a cell non-autonomous mechanism. Elife 10

24. Seale, P., Sabourin, L. A., Girgis-Gabardo, A., Mansouri, A., Gruss, P., and Rudnicki, M. A. (2000) Pax7 is required for the specification of myogenic satellite cells. Cell 102, 777–786

25. Yoshida, H., Matsui, T., Yamamoto, A., Okada, T., and Mori, K. (2001) XBP1 mRNA is induced by ATF6 and spliced by IRE1 in response to ER stress to produce a highly active transcription factor. Cell 107, 881–891

26. Kuang, S., Kuroda, K., Le Grand, F., and Rudnicki, M. A. (2007) Asymmetric self-renewal and commitment of satellite stem cells in muscle. Cell 129, 999–1010

27. Beauchamp, J. R., Heslop, L., Yu, D. S. W., Tajbakhsh, S., Kelly, R. G., Wernig, A., Buckingham, M. E., Partridge, T. A., and Zammit, P. S. (2000) Expression of CD34 and Myf5 defines the majority of quiescent adult skeletal muscle satellite cells. The Journal of cell biology 151, 1221–1234

28. Hindi, S. M., and Kumar, A. (2016) TRAF6 regulates satellite stem cell self-renewal and function during regenerative myogenesis. The Journal of clinical investigation 126, 151–168

29. Hollien, J., Lin, J. H., Li, H., Stevens, N., Walter, P., and Weissman, J. S. (2009) Regulated Ire1-dependent decay of messenger RNAs in mammalian cells. J Cell Biol 186, 323–331

30. Grandjean, J. M. D., Madhavan, A., Cech, L., Seguinot, B. O., Paxman, R. J., Smith, E., Scampavia, L., Powers, E. T., Cooley, C. B., and Plate, L. (2020) Pharmacologic IRE1/XBP1s activation confers targeted ER proteostasis reprogramming. Nature chemical biology 16, 1052–1061

31. Brack, A. S., Conboy, I. M., Conboy, M. J., Shen, J., and Rando, T. A. (2008) A temporal switch from notch to Wnt signaling in muscle stem cells is necessary for normal adult myogenesis. Cell stem cell 2, 50–59

32. Hayden, M. S., and Ghosh, S. (2004) Signaling to NF-κB. Genes & development 18, 2195–2224

33. Razani, B., Reichardt, A. D., and Cheng, G. (2011) Non-canonical NF-κB signaling activation and regulation: principles and perspectives. Immunological reviews 244, 44–54

34. Hindi, S. M., Shin, J., Gallot, Y. S., Straughn, A. R., Simionescu-Bankston, A., Hindi, L., Xiong, G., Friedland, R. P., and Kumar, A. (2017) MyD88 promotes myoblast fusion in a cell-autonomous manner. Nat Commun 8, 1624

35. Gardner, B. M., and Walter, P. (2011) Unfolded proteins are Ire1-activating ligands that directly induce the unfolded protein response. Science 333, 1891–1894

36. Maurel, M., Chevet, E., Tavernier, J., and Gerlo, S. (2014) Getting RIDD of RNA: IRE1 in cell fate regulation. Trends in biochemical sciences 39, 245–254

37. Urano, F., Wang, X., Bertolotti, A., Zhang, Y., Chung, P., Harding, H. P., and Ron, D. (2000) Coupling of stress in the ER to activation of JNK protein kinases by transmembrane protein kinase IRE1. science 287, 664–666

38. Quwaider, D., Corchete, L. A., Martín-Izquierdo, M., Hernández-Sánchez, J. M., Rojas, E. A., Cardona-Benavides, I. J., García-Sanz, R., Herrero, A. B., and Gutiérrez, N. C. (2022) RNA sequencing identifies novel regulated IRE1-dependent decay targets that affect multiple myeloma survival and proliferation. Experimental Hematology & Oncology 11, 1–15

39. Wu, R., Zhang, Q.-H., Lu, Y.-J., Ren, K., and Yi, G.-H. (2015) Involvement of the IRE1α-XBP1 pathway and XBP1s-dependent transcriptional reprogramming in metabolic diseases. DNA and cell biology 34, 6–18

40. Park, S.-M., Kang, T.-I., and So, J.-S. (2021) Roles of XBP1s in transcriptional regulation of target genes. Biomedicines 9, 791

41. Chèneby, J., Gheorghe, M., Artufel, M., Mathelier, A., and Ballester, B. (2018) ReMap 2018: an updated atlas of regulatory regions from an integrative analysis of DNA-binding ChIP-seq experiments. Nucleic acids research 46, D267–D275

42. Chèneby, J., Ménétrier, Z., Mestdagh, M., Rosnet, T., Douida, A., Rhalloussi, W., Bergon, A., Lopez, F., and Ballester, B. (2020) ReMap 2020: a database of regulatory regions from an integrative analysis of Human and Arabidopsis DNA-binding sequencing experiments. Nucleic acids research 48, D180–D188

43. Hammal, F., de Langen, P., Bergon, A., Lopez, F., and Ballester, B. (2022) ReMap 2022: a database of Human, Mouse, Drosophila and Arabidopsis regulatory regions from an integrative analysis of DNA-binding sequencing experiments. Nucleic acids research 50, D316–D325

44. Griffon, A., Barbier, Q., Dalino, J., van Helden, J., Spicuglia, S., and Ballester, B. (2015) Integrative analysis of public ChIP-seq experiments reveals a complex multi-cell regulatory landscape. Nucleic acids research 43, e27–e27

45. Liu, L., Cai, J., Wang, H., Liang, X., Zhou, Q., Ding, C., Zhu, Y., Fu, T., Guo, Q., and Xu, Z. (2019) Coupling of COPII vesicle trafficking to nutrient availability by the IRE1α-XBP1s axis. Proceedings of the National Academy of Sciences 116, 11776–11785

46. Goh, Q., and Millay, D. P. (2017) Requirement of myomaker-mediated stem cell fusion for skeletal muscle hypertrophy. Elife 6

47. Hindi, S. M., Sato, S., Xiong, G., Bohnert, K. R., Gibb, A. A., Gallot, Y. S., McMillan, J. D., Hill, B. G., Uchida, S., and Kumar, A. (2018) TAK1 regulates skeletal muscle mass and mitochondrial function. JCI Insight 3

48. Hindi, S. M., and Millay, D. P. (2022) All for One and One for All: Regenerating Skeletal Muscle. Cold Spring Harb Perspect Biol 14

49. Hindi, S. M., Tajrishi, M. M., and Kumar, A. (2013) Signaling mechanisms in mammalian myoblast fusion. Sci Signal 6, re2

50. Acosta-Alvear, D., Zhou, Y., Blais, A., Tsikitis, M., Lents, N. H., Arias, C., Lennon, C. J., Kluger, Y., and Dynlacht, B. D. (2007) XBP1 controls diverse cell type-and condition-specific transcriptional regulatory networks. Molecular cell 27, 53–66

51. Lin, J., Liu, H., Fukumoto, T., Zundell, J., Yan, Q., Tang, C.-H. A., Wu, S., Zhou, W., Guo, D., and Karakashev, S. (2021) Targeting the IRE1α/XBP1s pathway suppresses CARM1-expressing ovarian cancer. Nature communications 12, 5321

52. He, S., Fu, T., Yu, Y., Liang, Q., Li, L., Liu, J., Zhang, X., Zhou, Q., Guo, Q., Xu, D., Chen, Y., Wang, X., Chen, Y., Liu, J., Gan, Z., and Liu, Y. (2021) IRE1alpha regulates skeletal muscle regeneration through Myostatin mRNA decay. The Journal of clinical investigation

53. Iwawaki, T., Akai, R., Yamanaka, S., and Kohno, K. (2009) Function of IRE1 alpha in the placenta is essential for placental development and embryonic viability. Proc Natl Acad Sci U S A 106, 16657–16662

54. Benjamini, Y., and Hochberg, Y. (1995) Controlling the false discovery rate: a practical and powerful approach to multiple testing. J R Stat Soc B Methodol 57, 289– 300

55. Zhou, Y., Zhou, B., Pache, L., Chang, M., Khodabakhshi, A. H., Tanaseichuk, O., Benner, C., and Chanda, S. K. (2019) Metascape provides a biologist-oriented resource for the analysis of systems-level datasets. Nat Commun 10, 1523

56. Gu, Z., and Hubschmann, D. (2022) Make Interactive Complex Heatmaps in R. Bioinformatics 38, 1460–1462

