## Supplemental Figures S1-S5 and Table S1-S3 for "The IRE1α/XBP1 signaling axis drives myoblast fusion in adult skeletal muscle"

By

**Aniket S. Joshi, Meiricris Tomaz da Silva, Anirban Roy, Tatiana E. Koike, Mingfu Wu, Micah B. Castillo, Preethi H. Gunaratne, Yu Liu, Takao Iwawaki, and Ashok Kumar**

**This file contains:**

1) Figures S1-S5
2) Supplemental Table S1-S3

### Supplemental Figures

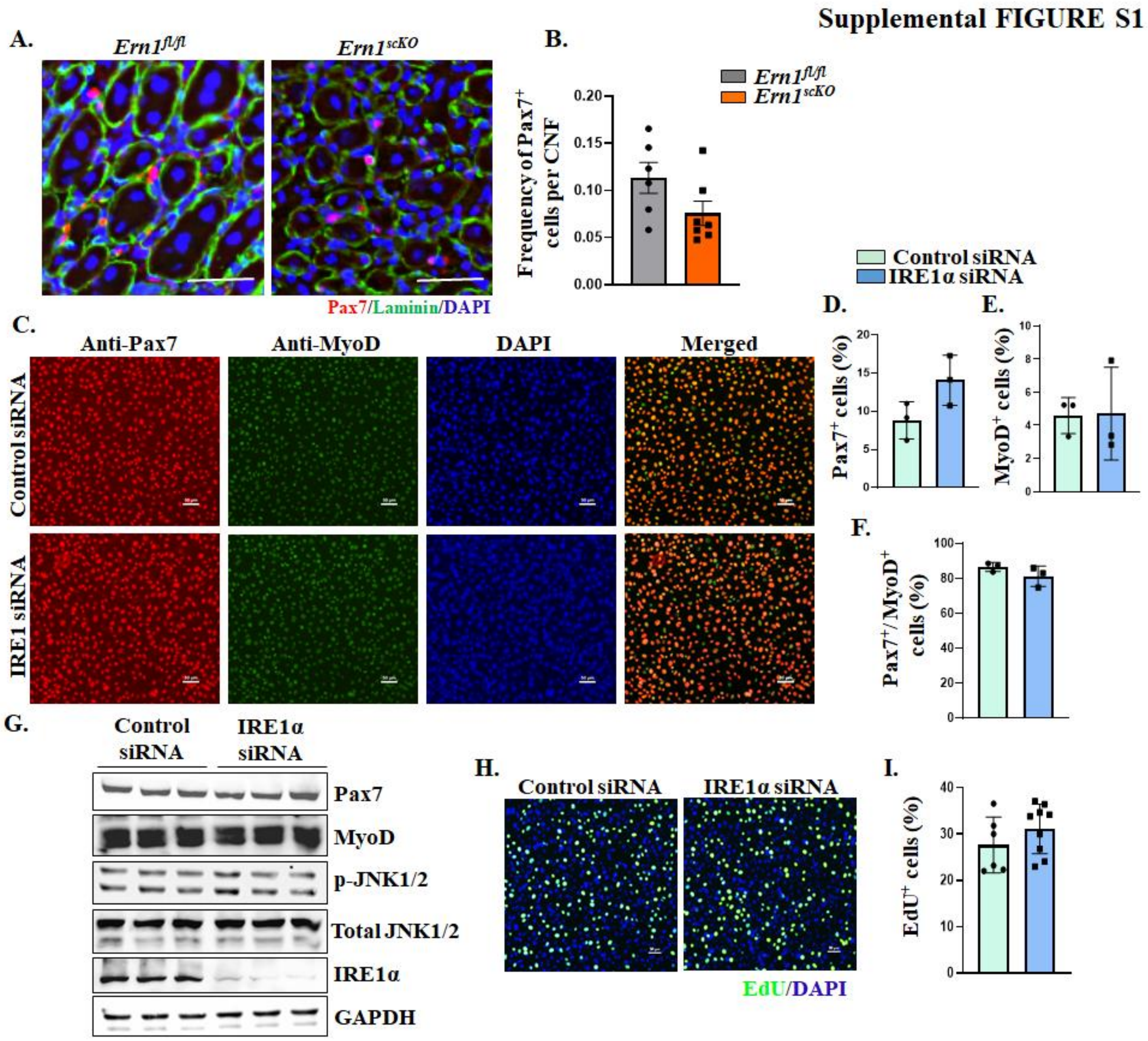


**Figure S1. Targeted ablation of IRE1α does not affect the abundance and proliferation of satellite cells. (A)** Representative images 5d-injured TA muscle of *Ern1*$^{fl/fl}$ and *Ern1*$^{scKO}$ mice after staining for Pax7 and Laminin protein. Nuclei was counterstained by DAPI. Scale bar: 50 µm. **(B)** Quantification of frequency of Pax7$^{+}$ cells per centrally nucleated myofiber (CNF) in 5d-injured TA muscle of *Ern1*$^{fl/fl}$ and *Ern1*$^{scKO}$ mice. n=6-7 per group. **(C)** Primary myoblasts were transfected with control or IRE1α siRNA for 24 h followed by immunostaining for Pax7 and MyoD protein. DAPI was used to stain nuclei. Representative photomicrographs are presented here. Scale bar: 50 µm. Quantitative analysis of proportion of **(D)** Pax7$^{+}$, **(E)** MyoD$^{+}$ and **(F)** Pax7$^{+}$/MyoD$^{+}$ cells in cultures transfected with control or

IRE1α siRNA. N=3 per group. **(G)** Western blot analysis for protein levels of Pax7, MyoD, p-JNK1/2, total JNK1/2, IRE1α, and an unrelated protein GAPDH in control and IRE1α siRNA transfected myoblast cultures. Primary myoblasts transfected with control or IRE1 siRNA were incubated with EdU for 1 h followed by detection of EdU in nuclei. DAPI was used to counterstain nuclei. **(H)** Representative photomicrographs of $EdU^+/DAPI^+$ control and IRE1α knockdown cultures. Scale bar: 50 µm. **(I)** Quantification of the proportion of $EdU^+$ cells in control and IRE1α knockdown myoblast cultures. n=6-9 per group. Data are presented as mean ± SEM.  No significant difference was observed by unpaired Student *t* test.

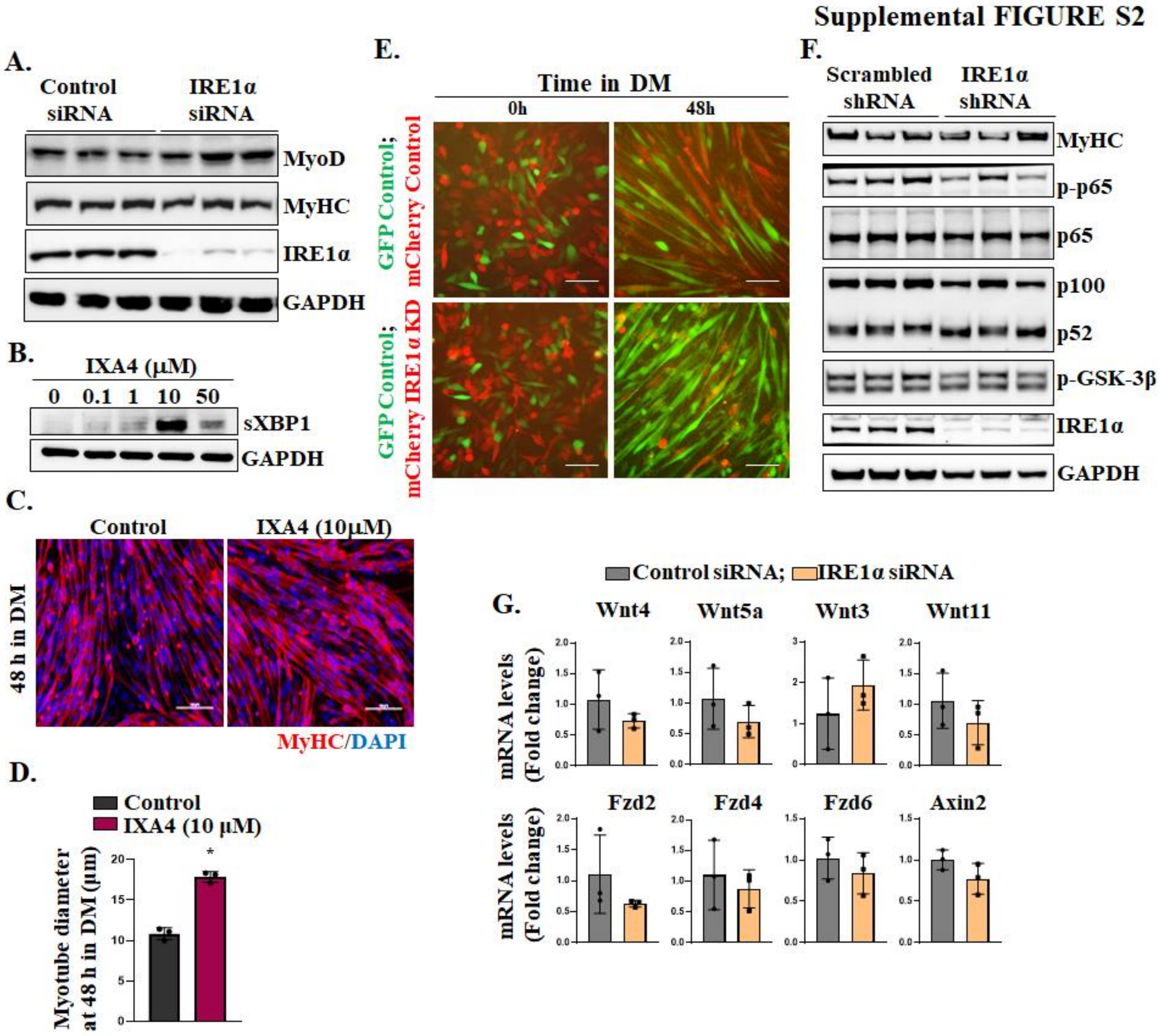


**Figure S2. Knockdown of IRE1α inhibits myoblast fusion without affecting NF-κB and Wnt pathways. (A)** Primary myoblasts were transfected with control or IRE1α siRNA for 24 h followed by protein extraction. Immunoblots presented here show levels of MyoD, MyHC, IRE1α, and GAPDH protein. **(B)** Primary myoblasts were incubated with different concentrations (0, 0.1, 1, 10, 50 µM) of IXA4 compound for 24 h and analyzed by performing western blot for sXBP1 and GAPDH protein. **(C)** Primary myoblasts were treated with vehicle alone (Control) or 10 µM IXA4 for 24 h followed by incubation in DM for 48 h. The myotube formation for evaluated by immunostaining for MyHC protein. DAPI staining was used to visualize nuclei. Representative photomicrographs are presented here. Scale bar: 50 µm. **(D)** Quantification of average myotube diameter in control and IXA4-treated cultures at 48 h of incubation with DM. Primary myoblasts were transduced with lentiviral particles expressing GFP protein (GFP Control), Scrambled shRNA (Control), or IRE1α shRNA (IRE1 KD) along with mCherry protein. Equal number of GFP Control and Control or IRE1α

KD cells were co-cultured followed by incubation in DM. **(E)** Representative photomicrographs of co-cultured myogenic cells expressing GFP and mCherry at 0 and 48 h of addition of DM. Scale bar: 50 μm. **(F)** Western blot analysis showing protein levels of MyHC, p-p65, p65, p100/p52, p-GSK-3β and IRE1α in scrambled or IRE1 shRNA expressing cultures at 24 h of differentiation. **(G)** Primary myoblasts were transfected with control or IRE1α siRNA for 24 h and analyzed for mRNA levels of *Wnt4a*, *Wnt5a*, *Wnt3a*, *Wnt11*, *Fzd2*, *Fzd4*, *Fzd6*, and *Axin2* using qRT-PCR analysis. n=3 per group. Data are presented as mean ± SEM. $^{*}p \leq 0.05$; values significantly different from control cultures analyzed by unpaired Student *t* test.

Supplemental FIGURE S3

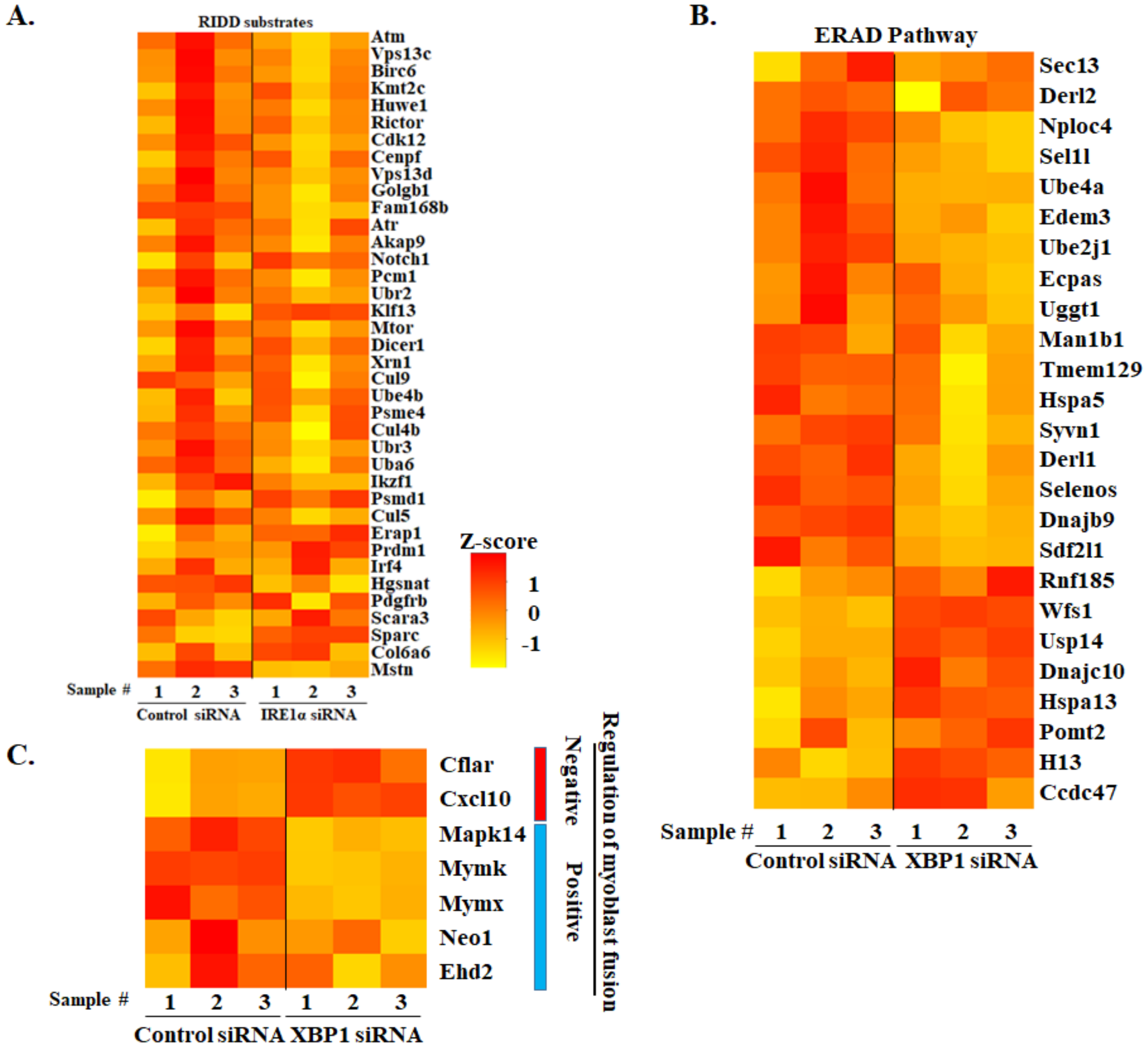


**Figure S3. Heatmaps depicting expression of various gene in IRE1α or XBP1 knockdown cultures. (A)** Heatmaps representing relative mRNA levels of selected putative RIDD substrates in IRE1α knockdown cultures. Heatmaps showing mRNA levels of **(B)** molecules associated with ERAD pathway, and **(C)** regulators of myoblast fusion in XBP1 knockdown cultures compared to control cultures at 24 h of incubation in DM.

Supplemental FIGURE S4

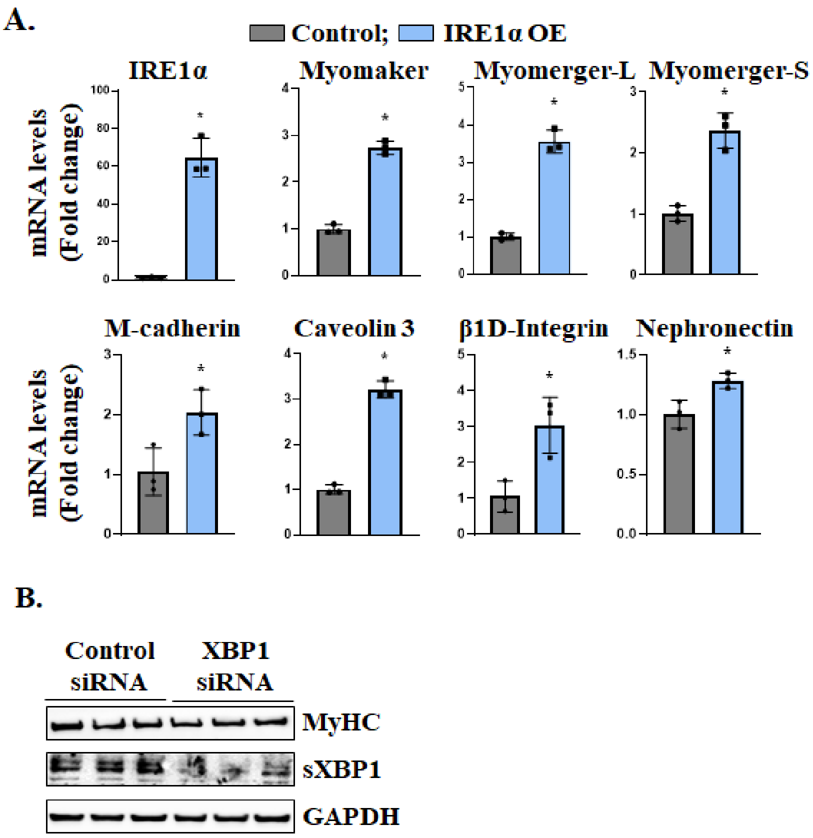


**Figure S4. Role of IRE1 on the gene expression of profusion molecules. (A)** Mouse myoblasts were transfected with control or XBP1 siRNA followed by incubation in DM for 24 h. Immunoblots presented here show the levels of MyHC, sXBP1, and GAPDH protein in control and XBP1 knockdown cultures. **(B)** Primary myoblasts were transduced with retrovirus expressing GFP (control) or IRE1α cDNA for 48 h. The cells were then used to measure mRNA levels of IRE1α, Myomaker, Myomerger-Long, Myomerger-Short, M-cadherin, Caveolin-3, β1D-Integrin, and Nephronectin by performing qRT-PCR assay. n=3 per group. Data are presented as mean ± SEM. $^*p \leq 0.05$, values significantly different from control cultures analyzed by unpaired Student *t* test.

### Supplemental FIGURE S5

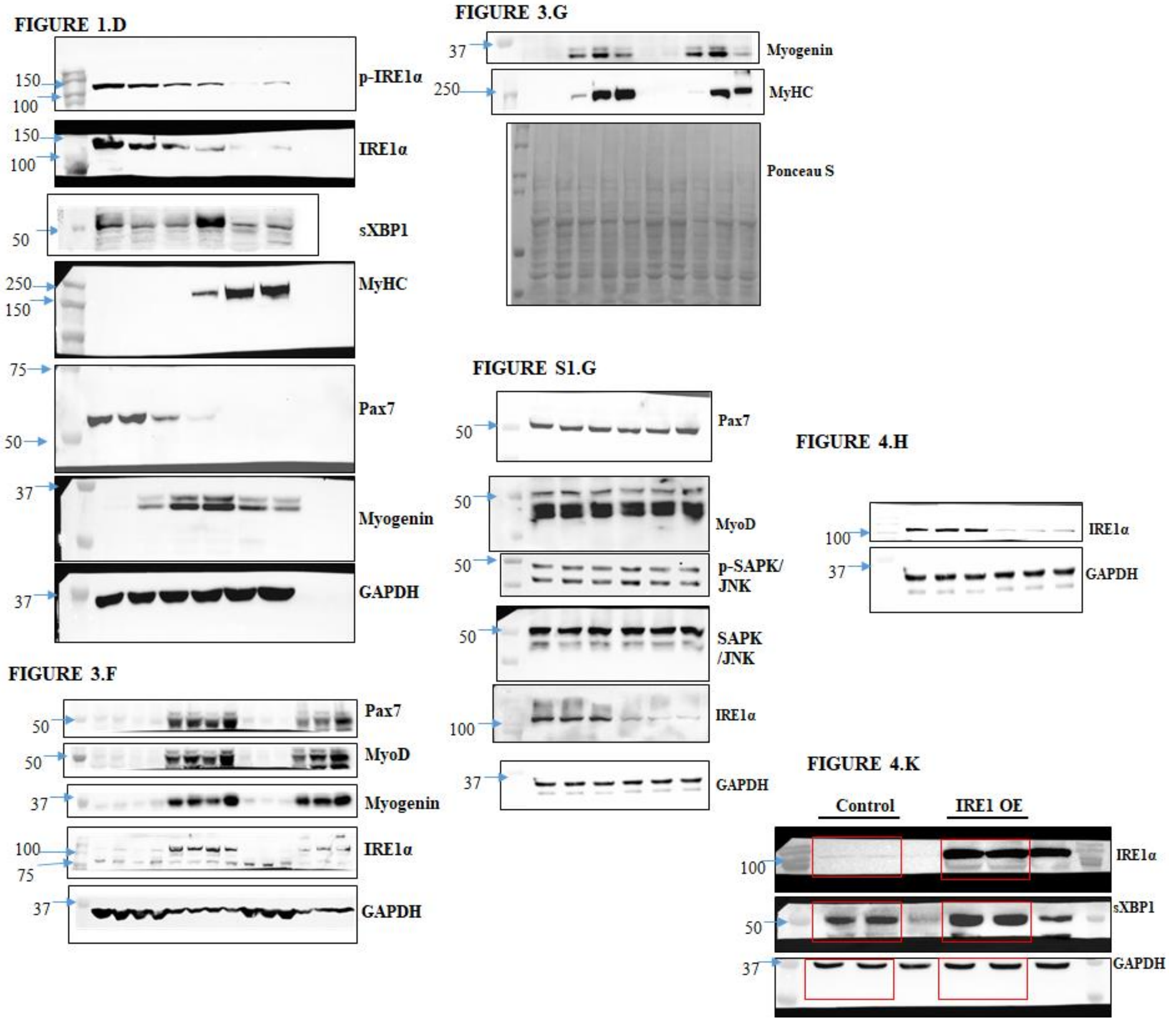

#### Supplemental FIGURE S5 (continued)

##### FIGURE 6.I

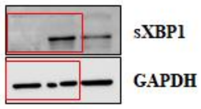


##### FIGURE S2.A

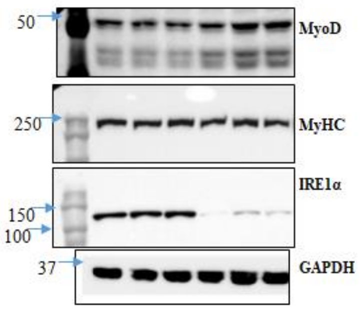


##### FIGURE S2.F

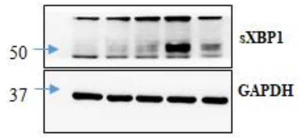


##### FIGURE S2.D

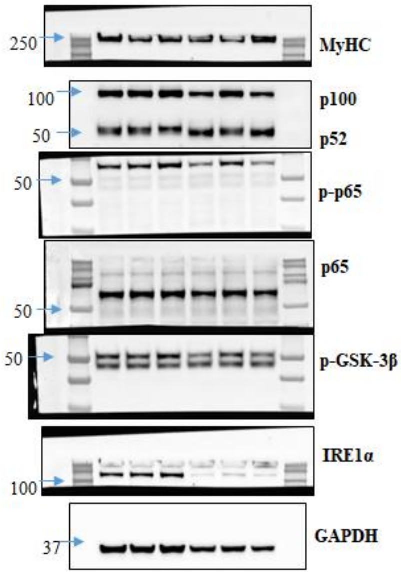


##### FIGURE 6.L

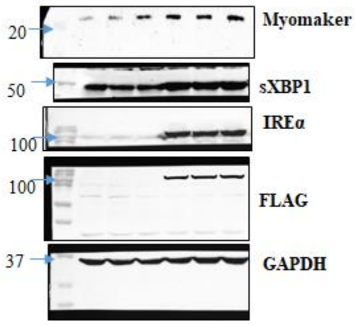


##### FIGURE 6.J

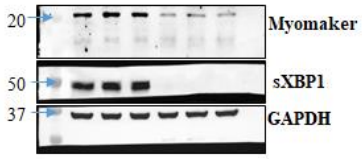


##### FIGURE 6.K

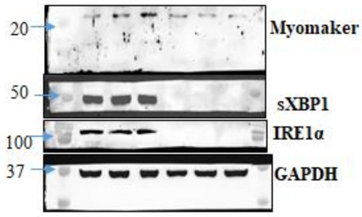


##### FIGURE S4.A

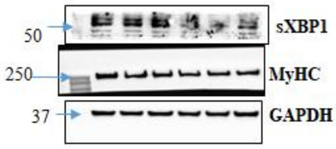


##### FIGURE 7.C

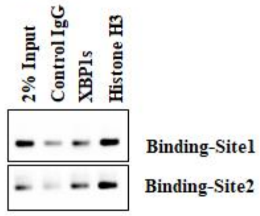

#### Supplemental FIGURE S5 (continued)

FIGURE 7.E

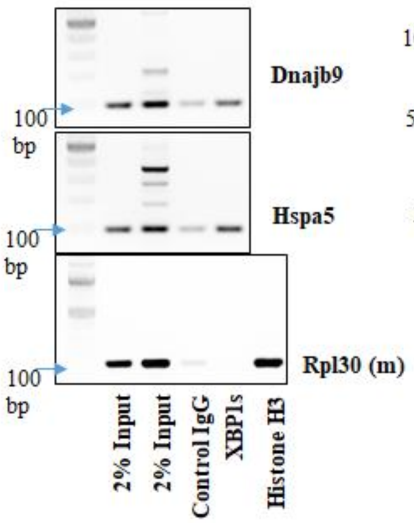


FIGURE 7.H

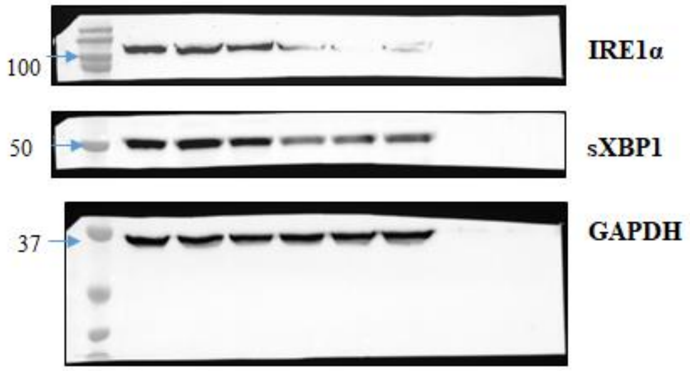


FIGURE 7.I

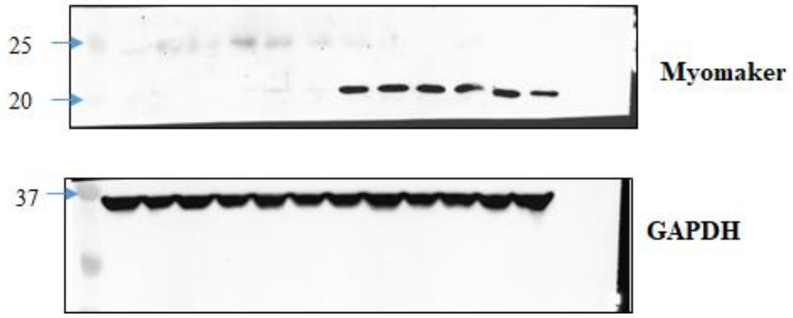


**Figure S5.** Uncropped gel images for western blot analysis and chromatin immunoprecipitation assays.

**Supplementary Table 1.** List of primers used for PCR/qRT-PCR analysis.

| Gene Name | Forward primer (5'-3') | Reverse primer (5'-3') |
|---|---|---|
| M-cadherin | TGGGCAGTCCCTGAGCCCAAA | TCCAGCGTGGCATTGAGGTACA |
| N-cadherin | CAGCAGATTTCAAGGTGGACGA | TCCTGGGTTTCTTTGTCTTGGG |
| BID-Integrin | CATCCCAATTGTAGCAGGCG | GAGACCAGCTTTACGTCCATAG |
| Caveolin-3 | GACCCCAAGAACATCAATGAGGAC | AGAAGGAGATACAGGCGAACAGGA |
| Myoferlin | CTACCAGAATGAGAATCGCTACCC | TACTCCCAGCCTTTCTCATCCA |
| ADAM12 | GGGCAAGAAGGCATAAGAGAGAGA | TGGTGAATGGGTCCTGGCTTAT |
| Myomaker | TATACTCCGGTCCCATAGGC | ATGCTCTTGTCGGGGTACAG |
| Myomerger-L | ACCAGCTTTCATGCCAGAAG | ATGTCTTGGGAGCTCAGTCG |
| Myomerger-S | CAGGAGGGCAAGAAGTTCAG | ATGTCTTGGGAGCTCAGTCG |
| IL-4 | GGATGTGCCAAACGTCCTC | GAGTTCTTCTTCAAGCATGGAG |
| IL-6 | CCTTCTTGGGACTGATGCTGG | GCCTCCGACTTGTGAAGTGGT |
| Nephronectin | CCAGAACAACTCCACTACCACCAA | CTGGGTCGTCCTTTACTTCCTCAT |
| Wnt3 | TGGGCCTGTCTTGGACAAA | GCGATGGCATGCACGAA |
| Wnt4 | CTGGAGAAGTGTGGCTGTGA | GGACGTCCACAAAGGACTGT |
| Wnt5a | GGCATCAAGGAATGCCAGTA | GTACGTGAAGGCCGTCTCTC |
| Wnt11 | GTAGGGCCTTCGCTGACAT | CGATGGTGTGACTGATGGTG |
| Fzd2 | CATCTCCATCCCGCTGTGCA | AGCACAGGAAGAAGCGCAGCTC |
| Fzd4 | GGCTACAACGTGACCAAGATGCC | GCACATTGGCACATAAACCGAAC |
| Fzd6 | GCGGCGTTTGCTTCGTT | CACAGAGGCAGAAGGACGAAGT |
| Axin-2 | TTTGGCACAGCTAGAGGAAG | TGGCTCTTTGTGATCTTCTGG |
| β-actin | CAGGCATTGCTGACAGGATG | TGCTGATCCACATCTGCTGG |
| Binding site 1 | GGTGACAGTCACGAAAGACC | GCCAGGTCCCTTTCCAAAT |
| Binding site 2 | CAGTTAAGGTCAAGCCTGGT | CACTCACCTTGTTCCTCCTC |
| Hspa5 | TGGTGGCATGGACCAATCAG | CGCCGACTCGCCTTATATAC |
| Dnajb9 | GAGCCGACCTACACGAAAC | AGGACCAAACGGCAACAA |

**Supplementary Table 2.** List of various antibodies used in the study.

| **Antibody** | **Source and Catalog no.** | **Dilution** | **Analysis** |
|---|---|---|---|
| Polyclonal rabbit-anti-phospho-ERN1 (S724) | Abnova # PAB12435 | 1:1000 / 1:200 | WB / IF |
| Monoclonal rabbit-anti-IRE1alpha | Cell Signaling Technology # 3294S | 1:1000 / 1:200 | WB / IF |
| Monoclonal rabbit-anti-XBP1-s (E9V3E) | Cell Signaling Technology # 40435 | 1:1000 / 1:50 | WB / ChIP |
| Monoclonal rabbit-anti-GAPDH | Cell Signaling Technology # 2118 | 1:1000 | WB |
| Monoclonal mouse-anti-FLAG-M1 | Sigma # F-3040 | 1:500 | WB |
| Monoclonal rabbit-anti-phospho-SAPK/JNK (T183/Y185) | Cell Signaling Technology # 4668S | 1:1000 | WB |
| Monoclonal rabbit-anti-SAPK/JNK | Cell Signaling Technology # 9252S | 1:1000 | WB |
| Monoclonal rabbit-anti-phospho-p65 NF-κB | Cell Signaling Technology # 3033 | 1:1000 | WB |
| Monoclonal rabbit-anti-total-p65 NF-κB | Cell Signaling Technology # 8242 | 1:1000 | WB |
| Polyclonal rabbit-anti-NF-κB p100/p52 | Cell Signaling Technology # 4882 | 1:1000 | WB |
| Polyclonal rabbit-anti-phospho-Glycogen synthase kinase-3 (GSK-3) | Cell Signaling Technology # 9336 | 1:1000 | WB |
| Monoclonal mouse-anti-Pax7 | DSHB # PAX7 | 1:200 / 1:100 | WB / IF |
| Monoclonal mouse-anti-Myosin heavy chain (embryonic) | DSHB # F1.652 | 1:200 / 1:15 | WB / IF |

| | | | |
|---|---|---|---|
| Monoclonal mouse-anti-MyoD | Santa Cruz Biotechnology # 377460 | 1:200 | WB / IF |
| Monoclonal mouse-anti-Myogenin | DSHB # F5D | 1:200 / 1:100 | WB / IF |
| Monoclonal mouse-anti-Myosin heavy chain | DSHB # MF20 | 1:250 / 1:100 | WB / IF |
| Polyclonal rabbit-anti-Laminin | Sigma # L9393 | 1:500 | IF |
| Goat anti-Mouse IgG1 Alexa Fluor 568 | Life Technologies # A21124 | 1:1000 | IF |
| Goat anti-Mouse IgG2 Alexa Fluor 594 | Life Technologies # A211135 | 1:1000 | IF |
| Goat anti-Rabbit IgG Alexa Fluor 488 | Life Technologies # 11034 | 1:1000 | IF |
| Goat anti-Mouse IgG2b Alexa Fluor 488 | Life Technologies # A21141 | 1:1000 | IF |

**Supplementary Table 3.** Mean TPM values for expression of various genes in control cultures.

| Gene | Mean TPM values | Gene | Mean TPM values |
|---|---|---|---|
| Hspa5 | 929.948 | Uba6 | 11.305 |
| Sdf2l1 | 105.930 | Pcm1 | 38.873 |
| Selenos | 150.429 | Akap9 | 18.635 |
| H13 | 46.874 | Golgb1 | 29.634 |
| Tmem129 | 41.276 | Atm | 22.331 |
| Derl1 | 144.536 | Fam168b | 58.945 |
| Man1b1 | 29.816 | Cul4b | 42.882 |
| Ccdc47 | 72.318 | Cul9 | 2.797 |
| Pomt2 | 7.664 | Ikzf1 | 0.045 |
| Rnf185 | 45.056 | Psme4 | 98.860 |
| Derl2 | 15.596 | Cenpf | 2.526 |
| Wfs1 | 137.619 | Atr | 2.190 |
| Dnajb9 | 64.526 | Xrn1 | 4.485 |
| Usp14 | 52.249 | Rictor | 4.565 |
| Uggt1 | 26.879 | Kmt2c | 4.851 |
| Syvn1 | 36.732 | Ubr2 | 26.618 |
| Sec13 | 108.220 | Birc6 | 15.102 |
| Ecpas | 43.393 | Huwe1 | 34.763 |
| Ube4a | 25.775 | Vps13c | 5.553 |
| Sel1l | 49.417 | Vps13d | 9.412 |
| Edem3 | 91.547 | Mtor | 28.880 |
| Hspa13 | 19.262 | Dicer1 | 13.641 |
| Ube2j1 | 76.995 | Ube4b | 65.523 |
| Dnajc10 | 42.990 | Notch1 | 17.609 |
| Nploc4 | 93.187 | Irf4 | 0.011 |
| Tmem182 | 190.196 | Mstn | 33.855 |
| Cxcl10 | 69.998 | | |
| Cflar | 6.785 | | |
| Cdon | 103.104 | | |
| Neo1 | 24.070 | | |
| Mymk | 2789.945 | | |
| Mapk14 | 53.559 | | |
| Ehd2 | 41.089 | | |
| Mymx | 594.790 | | |
